# Multimodal single-cell profiling of intrahepatic cholangiocarcinoma defines hyperactivated Tregs as a potential therapeutic target

**DOI:** 10.1101/2022.03.06.483155

**Authors:** Giorgia Alvisi, Alberto Termanini, Cristiana Soldani, Federica Portale, Roberta Carriero, Karolina Pilipow, Michela Polidoro, Barbara Franceschini, Ines Malenica, Simone Puccio, Veronica Lise, Giovanni Galletti, Veronica Zanon, Federico Simone Colombo, Michele Tufano, Alessio Aghemo, Luca Di Tommaso, Clelia Peano, Javier Cibella, Matteo Iannacone, Rahul Roychoudhuri, Matteo Donadon, Guido Torzilli, Paolo Kunderfranco, Diletta Di Mitri, Enrico Lugli, Ana Lleo

**Affiliations:** Laboratory of Translational Immunology, IRCCS Humanitas Research Hospital, via Manzoni 56, 20089, Rozzano, Milan; Bioinformatics Unit, IRCCS Humanitas Research Hospital, via Manzoni 56, 20089, Rozzano, Milan; Laboratory of Hepatobiliary Immunopathology, IRCCS Humanitas Research Hospital, via Manzoni 56, 20089, Rozzano, Milan; Laboratory of Tumor Microenvironment, IRCCS Humanitas Research Hospital, via Manzoni 56, 20089, Rozzano, Milan; Flow Cytometry Core, IRCCS Humanitas Research Hospital, via Manzoni 56, 20089, Rozzano, Milan; Humanitas University, Department of Biomedical Sciences, Via Rita Levi Montalcini 4, 20090 Pieve Emanuele – Milan, Italy; Division of Internal Medicine and Hepatology, Department of Gastroenterology, IRCCS Humanitas Research Hospital, Rozzano, Milan, Italy; Department of Pathology, IRCCS Humanitas Research Hospital, via Manzoni 56, 20089, Rozzano, Milan; Genomic Unit, IRCCS Humanitas Research Hospital, via Manzoni 56, 20089, Rozzano, Milan, Italy; Institute of Genetic and Biomedical Research, UoS Milan, National Research Council, Rozzano, Milan, Italy; Human Technopole, Viale Rita Levi Montalcini 1, 20157, Milano, Italy; Division of Immunology, Transplantation and Infectious Diseases, IRCCS San Raffaele Scientific Institute, 20132 Milan, Italy; Vita-Salute San Raffaele University, 20132 Milan, Italy; Experimental Imaging Centre, IRCCS San Raffaele Scientific Institute, 20132 Milan, Italy; Department of Pathology, University of Cambridge, United Kingdom. CB2 3QP; Division of Hepatobiliary and General Surgery, IRCCS Humanitas Research Hospital, via Manzoni 56, 20089, Rozzano, Milan, Italy

## Abstract

The quality of the immune infiltrate of intrahepatic cholangiocarcinoma (iCCA), a rare, yet aggressive tumor of the biliary tract, remains poorly characterized, limiting development of successful immunotherapies. We used high-dimensional flow cytometry to characterise the T cell and myeloid compartments of iCCA comparing these with their tumor-free peritumoral and circulating counterparts. We found poor infiltration of putative tumor-specific CD39+ CD8+ T cells accompanied by abundant infiltration of hyperactivated CD4+ regulatory T cells (Tregs), whose frequency in relation to that of CD4+ CD69+ T cells and conventional type 2 dendritic cells was associated with poor prognosis. Single-cell RNA-sequencing identified an altered network of transcription factors in iCCA-infiltrating compared to peritumoral T cells, suggesting reduced effector functions by tumor-infiltrating CD8+ T cells and enhanced immunosuppression by CD4+ Tregs. Specifically, we found that expression of mesenchyme homeobox 1 (MEOX1) was highly enriched in tumor-infiltrating Tregs, and demonstrated that MEOX1 overexpression is sufficient to reprogram circulating precursors to acquire the transcriptional and epigenetic landscape of tumorinfiltrating Tregs. Interfering with hyperactivated Tregs should be thus explored to enhance antitumor immunity in iCCA.

## INTRODUCTION

Cholangiocarcinoma (CCA) is a rare cancer that originates from the bile duct epithelia and accounts for 3-5% of all gastrointestinal malignancies worldwide (Tariq et al., 2019). Depending on the anatomic site of origin, CCA is divided into intrahepatic (iCCA), perihilar (pCCA), and distal (dCCA) CCA, with iCCA being the less prevalent type. However, the incidence rate of iCCA is constantly increasing and its mortality rate is extremely high, due to the aggressive evolution of the disease and the lack of efficient diagnostic and therapeutic treatments (Banales et al., 2020). Late diagnosis highly compromises surgery, the only current potentially curative option, and even among the 10-30% patients eligible for resection at diagnosis, 50% recur within the first year. Moreover, iCCA is a highly chemoresistant tumor, and pharmacological therapies are generally unsuccessful, with a 5-year survival rate that has persisted below 10% since the 1980s. Novel therapies targeting tumor subtypes with genetic rearrangements have been introduced last year into clinical practice after obtaining promising results in clinical trials (Abou-Alfa et al., 2020a; Abou-Alfa et al., 2020b), but only benefit 13-14% of patients (Nakamura et al., 2015; Sia et al., 2013). Therefore, there is an urgent need to develop valid therapeutic alternatives for iCCA.

During the last decade, immunotherapies approaches targeting checkpoint receptors expressed by tumor infiltrating lymphocytes (TILs) have revolutionized cancer treatment, increasing the overall survival (OS) of patients with multiple cancers (Borghaei et al., 2015; Snyder et al., 2014). The superior responsiveness to anti-programmed-death (PD)-1 checkpoint inhibitors is thought to be mediated by the unleashed reactivity of clonally expanded CD8+ T cells towards the cognate tumor antigens (Gros et al., 2014; Tumeh et al., 2014), among which those that are less differentiated, or stem-like, are endowed with enhanced functionality and capability for long-term persistence (Brummelman et al., 2018; Lugli et al., 2020; Miller et al., 2019; Sade-Feldman et al., 2018; Siddiqui et al., 2019). In this regard, tumors with high mutational burden respond better to checkpoint-inhibitor therapy (Gubin et al., 2014). Accordingly, tumors with mismatch repair (MMR)-deficiency and consequently high DNA microsatellite instability (MSI) are highly responsive to anti-PD-1 (Le et al., 2015). In that context, the FDA approved the use of anti-PD-1 antibody in 2017 in CCA and other solid tumors with MSI or MMR-deficiency, which nevertheless benefits only a small proportion of iCCA patients (Kim et al., 2020; Piha-Paul et al., 2020).

The tumor microenvironment is infiltrated by a diverse population of immune cells, among which effector and cytotoxic T cells and natural killer (NK) cells mediate tumor immunosurveillance and whose increased abundance is generally associated with delayed progression. By contrast, inhibitory subpopulations, including CD4+ regulatory T cells (Treg) and tumor-associated myeloid cells, can counteract immune responses and favor tumor growth (Binnewies et al., 2018). Several studies have recently deciphered the architecture of the tumor microenvironment of several cancers, expanding our knowledge of its complex composition and revealing novel drug targets. Such analyses now begin to reveal that many immune lineages are not compartmetalised into discrete subcompartments but, rather, comprise a continuum of functional phenotypes.

Knowledge regarding the complexity of the immune system in iCCA is still limited. Overall, iCCA is poorly infiltrated by the immune system, and is generally referred to as a “cold” tumor. Immunohistochemical analysis initially revealed the preferential presence of CD8+ and CD4+ T cells in tumors and in peritumoral areas, respectively (Kasper et al., 2009). A subsequent study in a cohort of 306 individuals with biliary tract cancers revealed that the longer OS correlated positively with a higher tumor infiltration of total CD4+ TIL, but did not distinguish CD4+ T cell subtypes (Goeppert et al., 2013). A recent study using single cell RNA-sequencing (scRNA-seq) of 8 paired tumoral and peritumoral samples elucidated at least in part the quality of immune cells infiltrating iCCA, revealing features similar to those found in other tumors, including the presence of activated Tregs and T cells expressing inhibitory receptors. The study also found a diverse population of cancer-associated fibroblasts (CAFs), where a predominant subpopulation of CD146^+^ vascular CAFs expressed high levels of IL-6, which in turn promoted tumor progression via IL-6R signaling on malignant cells (Zhang et al., 2020b). However, high-resolution well-powered datasets defining the architecture of the immune system infiltrating iCCA, both at the transcriptional and population level, are largely missing, thereby possibly preventing the development of future immunotherapy approaches.

Here we show that T cells within iCCA undergo profound remodeling, revealing the absence of putative tumor-specific CD39+ CD8+T_RM_ cells (Duhen et al., 2018; Simoni et al., 2018) and cytotoxic T lymphocytes (CTLs), and the abundant infiltration of a diverse population of Tregs, predicted to engage multiple inhibitory pathways on T and myeloid cells in the tumor microenvironment, while receiving signals that in turn may support their hyperactivated phenotype. A transcription factor (TF) novel to T-cell biology, mesenchyme homeobox 1 (MEOX1), was at least in part responsible for the transcriptional and epigenetic features of Tregs found in iCCA. We thus provide a high-resolution atlas of the T cell and myeloid cell infiltrate in iCCA, predict functional interactions between cell types with divergent functions and suggest that interfering with the activated Treg program might be needed to enhance anti-tumor immunity in iCCA.

## Results

### High-dimensional flow cytometry defines the T cell and myeloid cell composition of human iCCA

We initially generated single-cell suspensions from the tumor and the adjacent tumor-free tissue (hereafter referred to as peritumor), and isolated peripheral blood mononuclear cells (PBMCs) from 20 patients who were eligible for surgery shortly after diagnosis (**Table S1**). In addition, we isolated circulating immune cells that abundantly infiltrate the liver parenchyma by organ perfusion (hereafter referred to as perfusate). We next profiled millions of single cells with 2 high-dimensional flow cytometry panels (Brummelman et al., 2019) encompassing markers of T cell memory and effector differentiation, activation, cytotoxicity and exhaustion, CD4+ Treg markers as well as markers capable to define subsets of myeloid cells (**Fig. 1A**, **Fig. S1A** and **Table S2**). CD3+ cells, identifying the bulk of T cells, and CD3– CD66b– cells, enriching mainly for myeloid cells (hereafter referred to as “myeloid” for simplicity), were further analysed by PhenoGraph (Levine et al., 2015), a computational algorithm capable of clustering single cells without bias according to their relative expression of antigens in the multidimensional space. In this way, we identified 7, 10 and 12 CD4+, CD8+ and myeloid clusters, respectively (**Fig. 1B**), whose profile of antigen expression is shown in the heatmaps in **Fig. S1A**.

**Figure 1.**
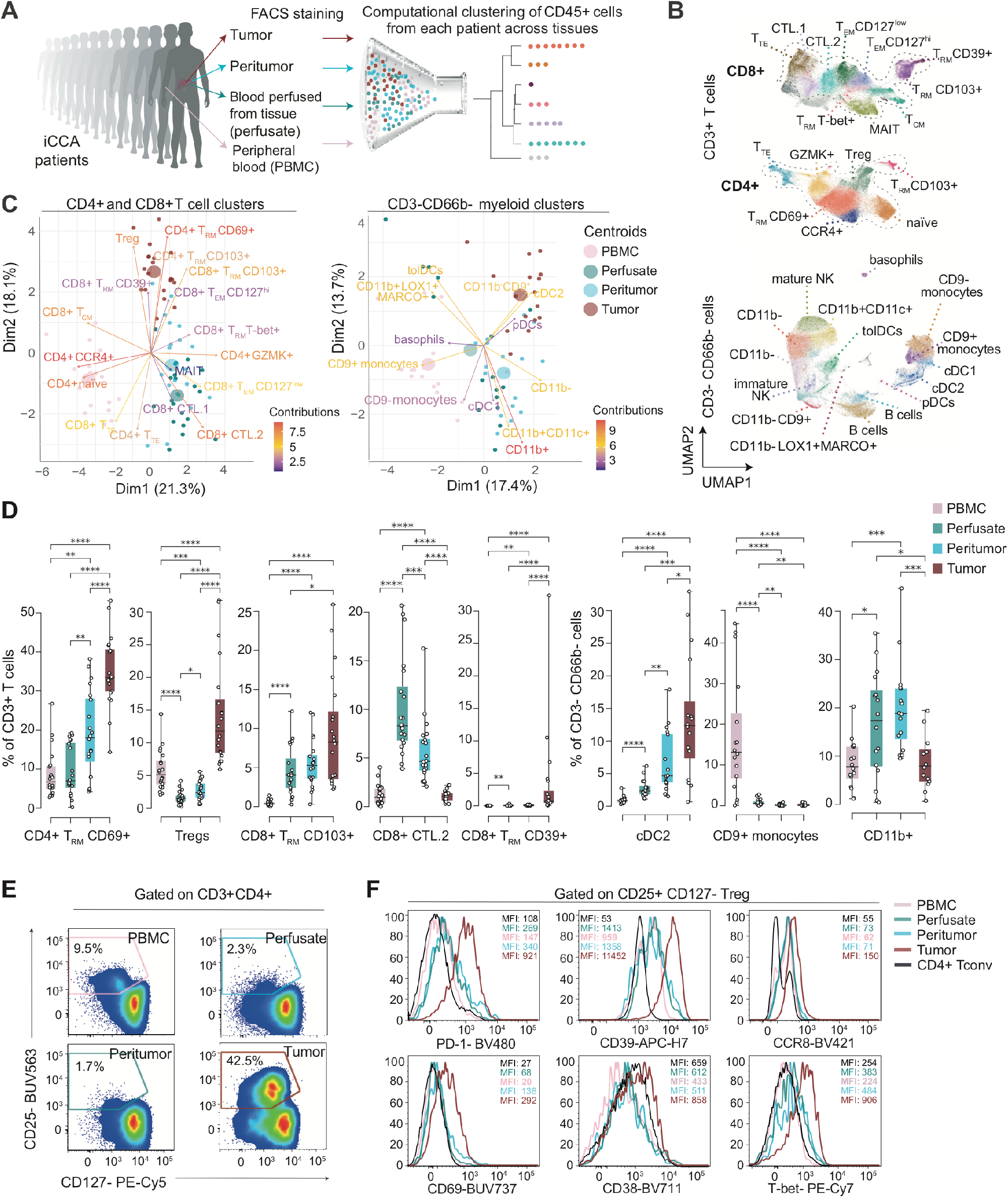
High-dimensional single-cell profiling of CD45+ cells infiltrating human intrahepatic cholangiocarcinoma (iCCA). **A.** Experimental workflow. **B.** UMAP representation of concatenated CD3+ T cell and CD3-CD66b-myeloid cell PhenoGraph clusters resulting from the flow cytometric analysis of paired tissue sites of iCCA patients (n=2O for T cell data; n=16 for myeloid data). Tregs: regulatory T; T_RM_: tissue-resident memory; T_CM_: central memory; T_EM_: effector memory; T_TE_: terminal effectors; MAIT: mucosal-associated invariant T; CTLs: cytotoxic T lymphocytes; NK: Natural Killer; cDC1: type 1 conventional dendritic cells; cDC2: type 2 conventional dendritic cells; pDCs: plasmacytoid dendritic cells; tolDCs: tolerogenic dendritic cells. **C.** PCA plots showing the distribution of samples according to the frequency of CD3+ T cell (left) and CD3-CD66b-myeloid cell (right) PhenoGraph clusters as in B. Small circles identify single samples, while big circles the mean of the distribution. Colors of the arrows and the cluster labels reflect the relative contribution to the PCA distribution. **D.** Box plots showing the median and the IQR of PhenoGrapg cluster frequency at different tissue sites. Bars indicated the SD. Dots depict single patient values. *=P<0.05, **=P<0.01, ***=P<0.001, ****=P<0.0001, two-sided Mann-Whitney test. **E.** Representative flow cytometric analysis of CD25 CD127-Tregs and **F.** of their markers from the different tissue sites. Conventional CD4+ CD25-CD127+ T cells (Tconv) are depicted as a control. Numbers indicate the percentage of cells in the gate (E) or the median fluorescence intensity (MFI) of marker expression.

Principal Component Analysis (PCA) of PhenoGraph cluster abundance revealed that the four tissue sites had a different T-cell and myeloid composition as a whole, although perfusate and the peritumor tended to share a similar immunophenotypic landscape (**Fig. 1C**). Specifically, PBMCs were characterized by the presence of CD4+ naïve and memory T cells expressing C-C chemokine receptor 4 (labelled as CD4+ CCR4+), CD45RO+ CCR7+ central memory CD8+ T cells (CD8+ T_CM_) as well as CD9+ and CD9– classical monocytes bearing a CD14+CD16– phenotype; perfusate and peritumor by CD4+ granzyme K+ CCR2^dull^ (CD4+ GZMK+) and, at a lesser extent, CD4+ GZMB+ 2B4+ terminal effector T cells (CD4+ T_TE_), by CD8+ T cells with a CD45RO-CCR7-GZMK+ GZMB+ phenotype (CD8+ CTL.2) or with a CD127^1ow^ CD45RO+ CCR7– GZMK+ GZMB+ effector memory T cell phenotype (CD8+ T_EM_ CD127^1ow^), by CD8+ T cells with a CD127^low^ GZMK+ CD161+ phenotype, labelled as MAIT cells (Dusseaux et al., 2011), and mainly by subsets of CD11b+ cells expressing or not CD11c, and of CD11b+CD11c+ HLA-DR^high^ CD141+ cells, suggesting the presence of immature myeloid cells and cDC1, respectively; tumors by subsets of CD4+ and CD8+ memory T cells expressing different combinations of the markers CD69 and CD103, collectively labelled as tissue-resident memory T cells (T_RM_; **Fig. 1C, D** and **Fig. S1B-D**). In the case of CD8+ T cells, a subset of CD69+ CD103+ T_RM_ also expressed high levels of CD39 (CD8 T_RM_ CD39+), a marker recently associated with CD8+ T cell reactivity to tumor antigens (**Fig. 1D**) (Duhen et al., 2018; Simoni et al., 2018). In line with their putative chronic stimulation by tumor antigens, these cells also expressed increased levels of the inhibitory receptor PD-1 and the activation marker CD38 compared to other iCCA-infiltrating CD8+ T cell clusters (**Fig. S1A**). Albeit present almost exclusively in tumors, the relative abundance of CD8+ CD39+ T_RM_ among total CD3+ was low (median=0.68, IQR: 0.41 and 3.32; **Fig. 1D**), possibly suggesting the poor immunogenicity of human cholangiocarcinoma or suppression of T cell responses against them. Notably, tumors were highly infiltrated by CD4+ CD127– CD25+ Tregs (**Fig. 1C-E**; **Fig. S1A**) overexpressing PD-1, CD39, CCR8, CD69, CD38 and the TF T-bet compared to Tregs from other tissue sites or the circulation (**Fig. 1F**), thereby indicating acquisition of a hyperactivated phenotype similar to that recently described in multiple other solid tumors (Alvisi et al., 2020; De Simone et al., 2016; Plitas et al., 2016). As far as myeloid cells were concerned, tumors were preferentially infiltrated compared to other tissue sites by CD11b+ CD11c+HLA-DR^high^ CD1c+ cDC2 (**Fig. 1D**), and by CD11b– CD9+ cells. Tumors were also infiltrated by CD11c– HLA-DR^high^ CD123+ plasmacytoid DCs (pDCs), which were also abundant in the perfusate and in the peritumoral area, less so in the PBMCs, and by CD11b+ CD11c+ immature myeloid cells, which instead were ubiquitous (**Fig. 1C-D**; **Fig. S1D**). Overall, these data indicate that human iCCA is characterized by a different landscape of T cell and, at a lesser extent, myeloid phenotypes.

### Impact of the immune landscape at surgery on the prognosis of iCCA patients

We next investigated whether a different immune landscape could influence disease progression in our cohort of 20 patients. While the relative abundance of single clusters alone was not informative in this regard (data now shown), combinatorial analysis of cluster abundance defined that the frequency of Tregs as relative to that of CD4+ CD69+ T_RM_ cells or as that of cDC2 were highly associated with disease free survival (DFS). Specifically, worse prognosis was associated with high Treg infiltration and either low CD4+ CD69+ T_RM_ cell or low cDC2 infiltration (**Fig. 2**). Interestingly, a recent report mechanistically related Treg-dependent inhibition of cDC2 activity with tumor immunosuppression using preclinical models (Binnewies et al., 2019). Accordingly, the relative abundance of these two cell types was associated with disease progression of head and neck squamous cell carcinoma and response of melanoma to checkpoint blockade. Overall, these data suggest that a common axis regulates anti-tumor immunity in different cancer types, and highlight the important role that CD4+ Tregs might play in the disease course of iCCA.

**Figure 2.**
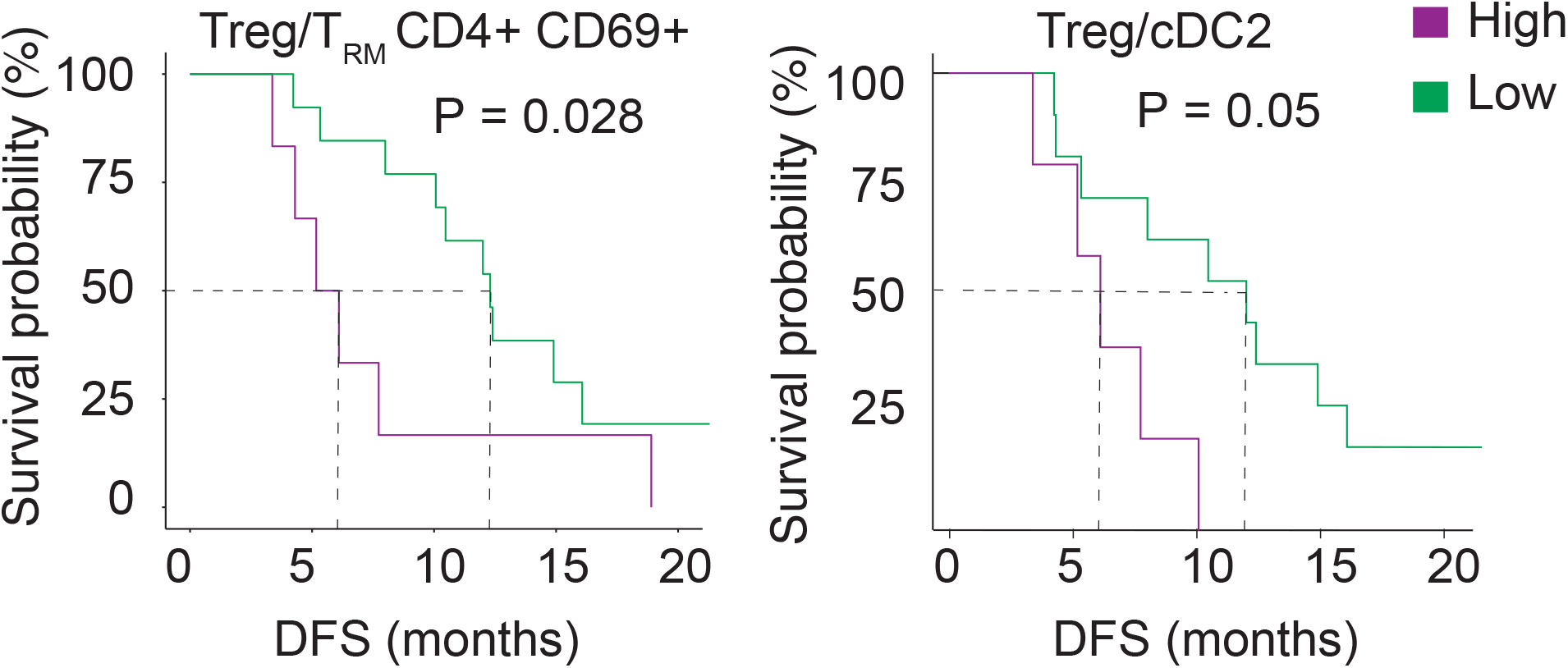
Features of immune infiltrate that predict disease-free survival (DFS) in iCCA. Kaplan-Meier DFS curves according to the intra-tumoral frequencies of Tregs as related to that of T_RM_ CD4+ CD69+ cells (left) or cDC2 (right) in each patient (n = 16). The cohort was subdivided in 2 groups according to the percentile rank (set at 0.7). The p value (P) was calculated by Gehan Breslow-Wil-coxon test.

### scRNA-seq reveals tumor-specific differences in gene expression by specific T cell subpopulations

We next performed scRNA-seq of CD45+ immune cells isolated from 6 cholangiocarcinomas and paired peritumoral tissues to gain more insights on the molecular characteristics of the tumor-specific immune infiltrate. Cluster analysis of scRNA-seq data and subsequent enrichment of defined immune signatures revealed that, among immune cells, T and NK cells were most abundant, followed by myeloid cells and B cells (**Fig. 3B** and **Fig. S2A,B**). CD45– stromal/tumor cells from 2 patients, spiked in at known concentration as a control, separated from these subsets of immune cells (**Fig. 3A**). Among CD45+ cells, tumors tended to harbor increased frequencies of T cells, reduced frequencies of NK and B cells, and similar frequency of myeloid cells compared to adjacent peritumoral tissue (**Fig. S2B**). Overall, tumoral and peritumoral tissues could be clearly distinguished on the basis of T and NK cell gene expression profiles, and less so by B cell and myeloid profiles, indicating that T cells and NK cells undergo specific transcriptional changes in the tumor (**Fig. S2C**). Inspired by flow cytometry results, we focused our subsequent investigation on T cells and myeloid cells, and subclustered these populations of cells to further identify their transcriptional characteristics within tumors compared to the adjacent peritumoral tissue. As scRNA-seq datasets may be characterized by zeros and dropout events, we employed imputation of single-cell data, a computational approach capable to infer gene expression even in the presence of dropouts, to improve the detection of gene expression (Linderman et al., 2018). For instance, this approach improved the detection of *CD4* and *CD8A* mRNA expression (**Fig. S3A**). We identified 8 clusters of myeloid and 11 clusters of T cells, reflecting, at least in part, those subpopulations that were identified by flow cytometry. Among myeloid, we identified *CD14*^high^ classical monocytes (C0) expressing *S100A8*, *VCAN* and *CD36*, among other genes; *ID3*^high^ macrophages (C1) expressing *VSIG4* and resembling tissue resident Kupffer cells, as previously suggested (Mass et al., 2016); *MARCO*^high^ macrophages (C3) expressing *PLIN2*, *APOC1* and *SPP1*, among others; *TREM2*^high^ macrophages (C5) expressing *APOC1* and *C1QA/B/C;* cDC2 (C2), expressing *FCER1A*, *ADAM8*, *CD1C*, *CLEC10A* and *IRF4*; cDC1 (C7), expressing *IRF8*, *IDO1*, *CLEC9A* and *BATF3*; and non-classical monocytes (C6) expressing *FCGR3A*, *CDKN1C*, *LILRA1* and *LILRB2* (Zhang et al., 2020a). Among T cells, we identified subsets of early differentiated memory T cells (C2, C5 and C8), expressing different combinations of genes previously related to stem-like memory cell differentiation such as *CCR7*, *GPR183*, *IL6R*, *SATB1*, *CCR4* and *IL7R* (Galletti et al., 2020); subsets of effector cells (C0, C3, C6 and C7), expressing different combinations of genes previously related to effector memory differentiation, such as *CSF1* (encoding M-CSF), *TNF*, *IFNG*, *TBX21* [encoding the TF T-bet], *ID2*, *PRDM1* (encoding the TF BLIMP-1), *GZMA*, *GZMK* and the C-C chemokines *CCL4* and *CCL5*, among others; a terminally differentiated/cytotoxic subset (C1), expressing the cytotoxicity-related genes *CRTAM*, *NKG7* and *PRF1* (encoding perforin) and the terminal differentiation-related TF *ZEB2*; a T_RM_ cell subset (C4), expressing *ITGAE* (encoding CD103 integrin), *CISH*, *CCR2*, *HOPX* as well as detectable levels of *ENTPD1* (encoding CD39), suggesting potential tumor reactivity (Duhen et al., 2018; Simoni et al., 2018) (**Fig. 3D and Fig. S3B**); and a subset expressing several CD4+ Treg-related genes (C10), including *IL2RA*, *FOXP3*, *BATF*, *TIGIT*, *CD177*, *IL1R2*, among others (**Fig. 3D**). An additional subset, C9, was found to express *CCL20*, *IL23R*, *RORC* and *KLRB1* (encoding CD161), suggesting the identification of CD8+ mucosal associated invariant T (MAIT) cells (Dusseaux et al., 2011) or, alternatively, CD4+ T helper type-17 cells (Th17). The poor expression of *CD4* and *CD8A* by C9 (**Fig. S3B**) precluded further distinction between these two subsets.

**Figure 3.**
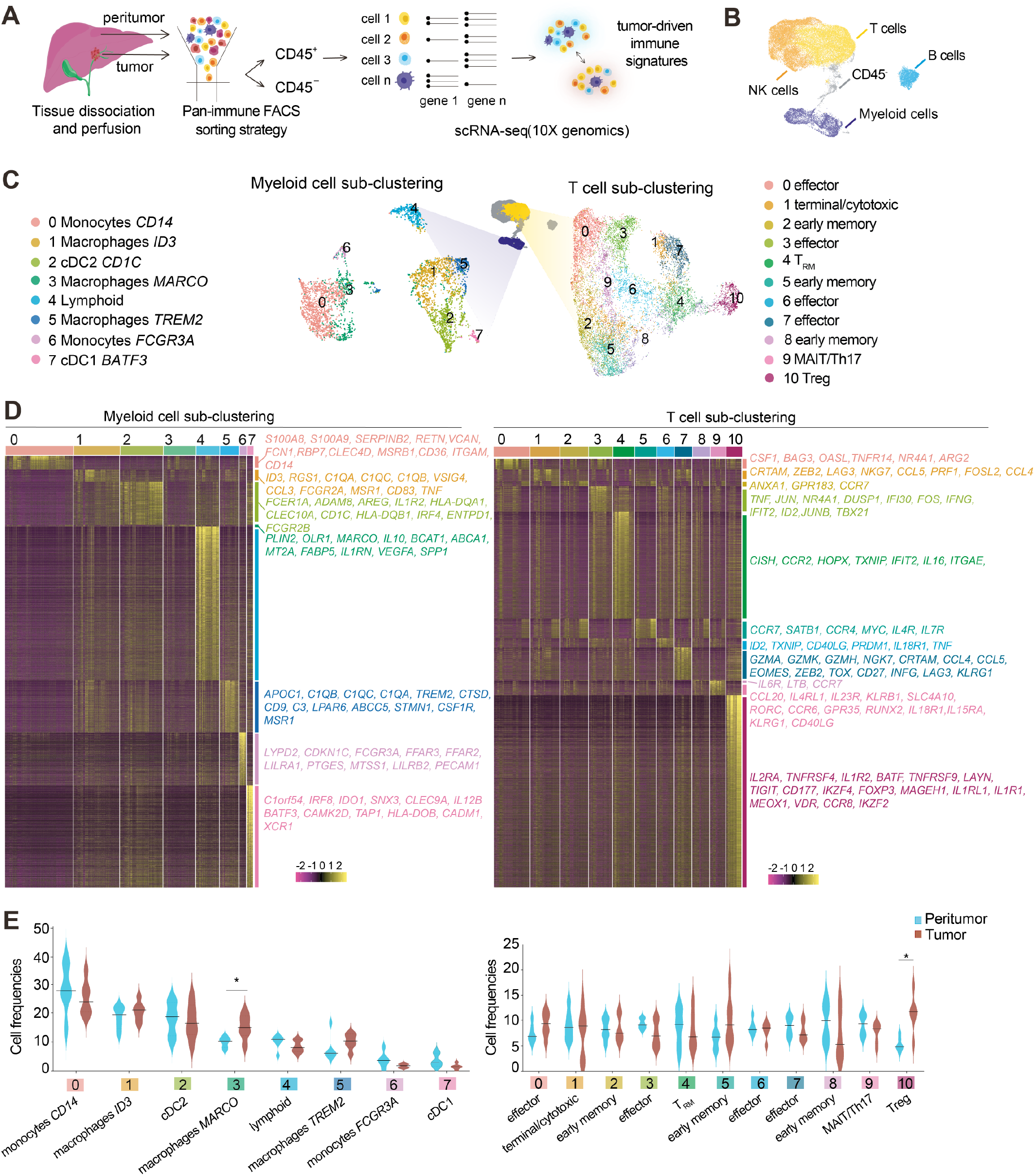
Single cell RNA-sequencing (scRNA-seq) reveals tumor-specific differences in gene expression by specific T-cell populations. **A.** Workflow of the experimental strategy: scRNA-seq was performed on paired peritumoral and tumoral surgical resections from iCCA patients (n = 6). **B**. UMAP projection of all cells analyzed (n = 31,745). Each dot corresponds to one single cell, colored according to immune cell lineage. **C**. UMAP projection representing T (n = 12,644) and myeloid cells (n = 4,585) sub-clustering. Clusters are depicted with different colors. T_RM_: tissue-Resident Memory T cells; Tregs: regulatory T cells; cDC2: type 2 dendritic cells; cDC1: type 1 dendritic cells. **D.** Gene expression heatmap of myeloid and T cell cluster marker genes after Adaptively-threshold Low-Rank Approximation (ALRA) imputation. Columns: cells grouped by clusters. Rows: cluster marker genes after filtering by Fold Change (|log_2_(FC)|>0.5), q-value<0.05 and percentage of expression in cluster cells (pct1)>0.5. Representative genes that are differentially expressed in specific clusters are labelled on the right. **E**. Violin plots showing the relative distribution of each T and myeloid cell cluster among tumoral (dark red) and peritumoral (light blue) samples. Lines represent median frequencies between samples from 6 patients, *p<0.05 with 2-way ANOVA test.

Overall, scRNA-seq revealed increased abundance of *MARCO^high^* myeloid cells and of CD4+ Tregs in tumoral compared to peritumoral tissues (**Fig. 3E**), in line with results obtained by flow cytometry, although could not detect additional differences, likely due to the limited number of patients that could be analysed using scRNA-seq. Nevertheless, scRNA-seq identified differences in T-cell gene expression between these two sites (**Fig. S3B**), suggesting distinct functional regulation of T cells in tumors. The biggest differences were observed among Tregs, where those from tumors expressed increased levels of *CTLA4*, *HAVCR2* (encoding the inhibitory receptor TIM-3), *TIGIT*, *BTLA* and *ENTPD1* (**Fig. S3B**), among others (**Fig. S3D**), confirming previous flow cytometry data. By contrast, C4 T_RM_ and C7 effector subsets from tumors tended to express increased levels of *HAVCR2* and *CTLA4* (only C7) and lower levels of the effector/killer molecules compared to those from the peritumoral tissue (**Fig. S3C**). Overall, these data suggest functional modulation of the T-cell infiltrate in the tumor microenvironment, with heightened activation of Tregs and reduced functional capacity of putative tumor-specific *ENTPD1*^high^ T_RM_ cells and effector T cells.

### Dynamic remodeling of the Treg cell interactome in iCCA

Our high-dimensional single-cell profiling identified transcriptional and proteomic modulation of Treg ligands and receptors in tumoral vs. peritumoral tissues, potentially implicating that Tregs differentially interact with the surrounding microenvironment. To gain more insights into the molecular identity of these interactions, we used CellPhoneDB, a computational algorithm capable of predicting cell-cell communications from differentially expressed ligand:receptor (L:R) pairs in single-cell data (Vento-Tormo et al., 2018). We found that, overall, Tregs interacted with different T cell and myeloid subsets via multiple interactions that were more significant in tumoral than in peritumoral tissues. The repertoire of these interactions, that involved co-inhibitory and co-stimulatory signals, TNF superfamily members, cytokines, chemokines and their receptors, tended to be different among T-cell clusters, while relatively uniform among myeloid clusters (**Fig. 4A, B**). Among others, we found enhanced interaction between CD80 and CD86 expressed by myeloid cells and CD28 costimulatory receptor expressed by Tregs. Similarly, the TNF superfamily members CD70 and TNFSF4 (encoding OX-40 ligand), overexpressed by T cell subsets, and TNFSF9, encoding 4-1BB ligand and expressed by both T and myeloid cells, interacted with their cognate receptors CD27, TNFRSF4 (OX-40) and TNFRSF9 (4-1BB) in tumors, suggesting that these interactions are important for the maintenance of activated Tregs as recently shown in murine models (Vasanthakumar et al., 2017). Importantly, CD80 and CD86 also showed enhanced predicted interaction with CTLA4, highly expressed by Tregs in tumors and important for the Treg-mediated inhibition of immune responses via competition with CD28. Additional interactions of note that were stronger in tumoral than in peritumoral tissues involved ligands expressed by myeloid subsets such as ICOSLG (encoding ICOS ligand), mediating Treg proliferation and functional stability following interaction with the cognate receptor ICOS (Kornete et al., 2012), or NECTIN2/NECTIN3/PVR and PDCD1LG2 (encoding PD-L2) on myeloid cells interacting with the inhibitory receptors TIGIT and PDCD1 (encoding PD-1), respectively. Moreover, intratumoral Tregs preferentially expressed CD200 and SIRPG inhibitory ligands interacting on myeloid cells with CD200R1 and CD47, respectively, both possibly involved in the downregulation of inflammatory responses (Vaine and Soberman, 2014; Weiskopf, 2017). Additional interactions involved members of the IL-1 family, potentially suggesting a role in Treg biology in tumors that will need further investigation. Thus, iCCA-infiltrating Tregs are characterized by extensive remodeling of the expression of receptor ligand pairs required for cell-cell communication, suggestive of enhanced Treg-mediated immunosuppression in the iCCA microenvironment.

**Figure 4.**
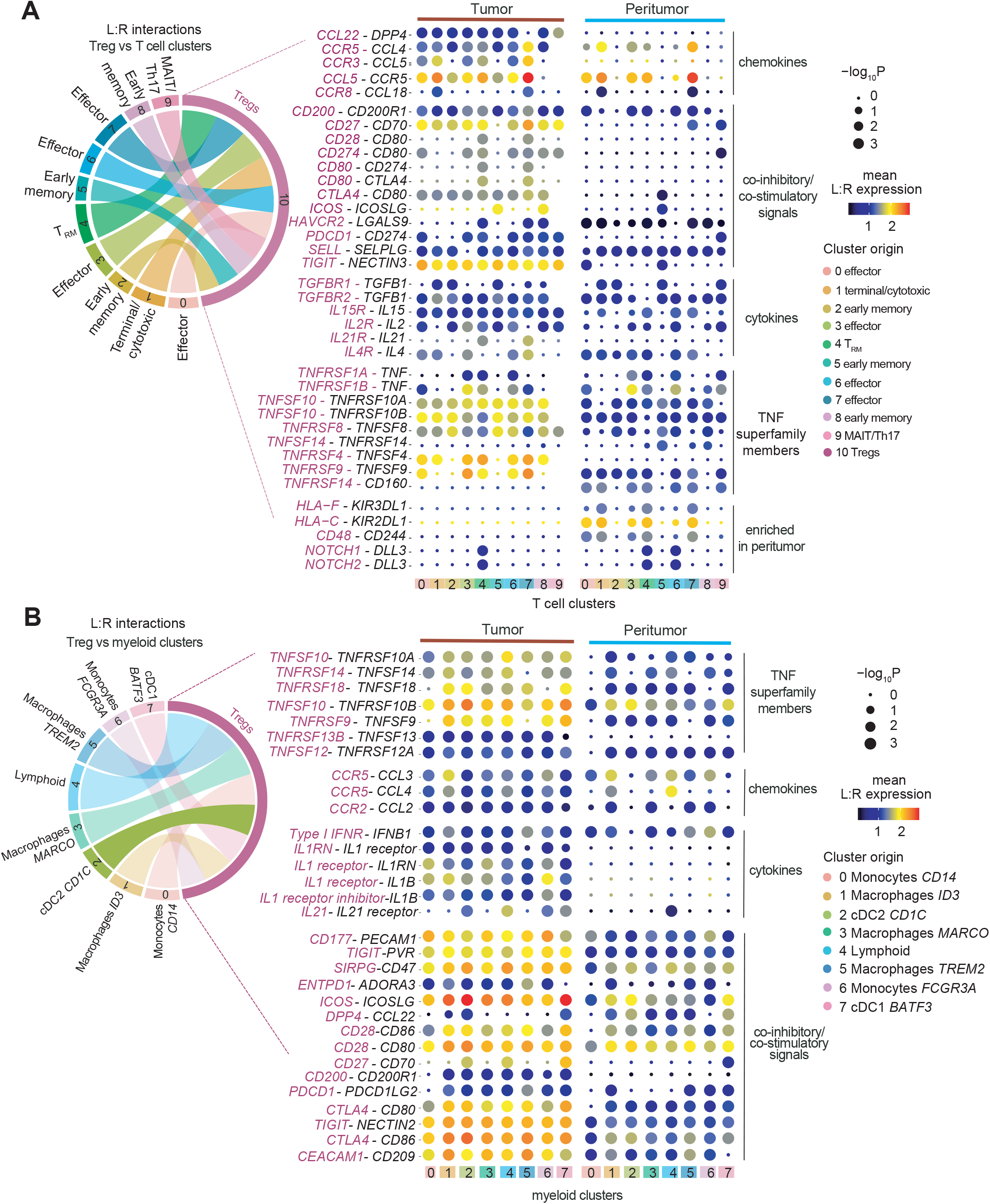
Dynamic remodeling of the Treg cell interactome in iCCA. CellPhoneDB intercellular communication analysis between Tregs (scRNA-seq cluster 10) and (**A**) T cell or (**B**) myeloid cell clusters identified by scRNA-seq. In both A and B, circos plot show all predicted cell–cell interaction events via ligand:receptor (L:R) pairs, while bubble plots indicate the mean L:R expression (color scale) and the corresponding P value (size of the bubble). Molecules expressed by Tregs and the interacting populations are in purple and in black, respectively. P: empirical permutation P value.

### Transcriptional network inference to understand the molecular basis of diminished effector T cell activation and enhanced Treg activation in iCCA

TF control the expression of several genes simultaneously and their co-regulation defines cell fate and functional responses. We hypothesized that differences in TF activities could be at the basis of differences in tumoral vs. peritumoral T-cell gene expression. To test this, we applied SCENIC, a computational algorithm capable to predict TF activity by the analysis of TF motifs that are enriched at the promoters of expressed genes in our scRNA-seq data (Aibar et al., 2017) (**Fig. 5A**). SCENIC analysis clearly separated tumoral and peritumoral T cells (**Fig. 5A**) as well as the majority of T-cell clusters previously defined by scRNA-seq (**Fig. 5B**), thus indicating that tissue-derived T-cell states can be described by their inferred TF activity. Among others, we found that IRF2, IRF3 and STAT1-mediated transcriptional activities, possibly dependent on type I interferon signaling and involved in promoting effector functional capacity (Huber and Farrar, 2011), were reduced in tumoral vs. peritumoral T cell clusters, especially in C3, C6 and C7 of effector cells and, at a lesser extent, in C4 of T_RM_ cells (**Fig. 5B**). C7 along with C1 of terminal/cytotoxic T cells from tumors vs. peritumors also showed reduced transcriptional activities of RUNX3, EOMES and TBX21 TFs, which mediate the expression of effector and cytotoxic molecules (Cruz-Guilloty et al., 2009). Altogether, these data suggest loss of T_RM_ and effector T-cell functionality in cholangiocarcinoma is due to altered, cell type-specific transcriptional programs. By contrast, tumor-infiltrating Tregs displayed increased activity of several TFs compared to those infiltrating the peritumor, including FOXP3, the lineage-specification factor for Treg cells, IKZF2, linked to stability of the Treg lineage (Getnet et al., 2010), IRF4 and its transcriptional partner BATF, recently reported to play a pivotal role in Treg activation and suppression in tumors (Alvisi et al., 2020), STAT5A and SMAD1, possibly reflecting IL-2 and TGF-β signaling, respectively, FOXA1, linked to enhanced suppressive function (Liu et al., 2014), and several others, such as VDR, SOX9, ZEB1 and Mesenchyme Homeobox 1 (MEOX1), whose functions in Treg biology remain poorly described or unknown (**Fig. 5B**).

**Figure 5.**
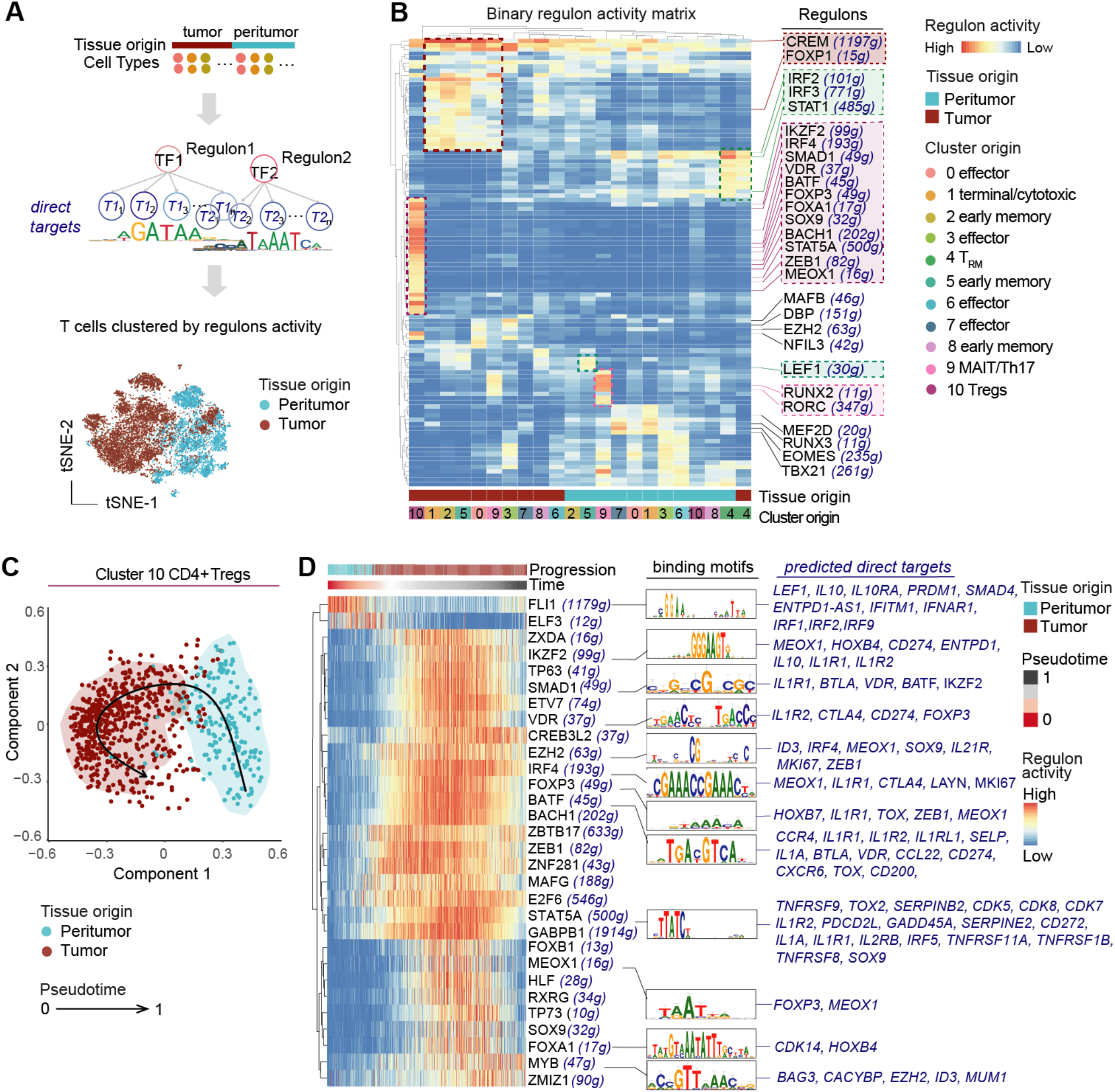
Transcriptional network inference to understand the molecular basis of diminished effector T cell activation and enhanced Treg activation in iCCA. **A.** Top: schematic overview of the SCENIC pipeline. Bottom: t-SNE map of all T cells based on regulon activity scores, as obtained by SCENIC. **B.** Heatmap of binary regulon activity (arbitrary threshold = 0.5) by scRNA-seq T cell clusters related to peritumoral (light blue) and tumoral (dark red) tissues. Relevant transcription factors (TFs) are indicated. The putative number of genes regulated by each TF is in brackets. Colored dashed boxes in the heatmap identify the T cell cluster with the highest TF activities. **C.** Zoom-in view of the scRNAseq cluster 10 (CD4+ Tregs) regulatory state. Tregs are distributed in a trajectory map by SCORPIUS according to regulon activity scores and color-coded based on tissue origin. **D.** Pseudo-time alignment of regulon activity in single Tregs determined by SCORPIUS trajectory analysis. The top 30 TF leading the time line are indicated along with their DNA binding motifs and their predicted targets (manually selected; in dark blue).

We next ordered TF activities in pseudotime by using a dedicated algorithm, i.e., SCORPIUS (Saelens et al., 2019), so to possibly identify specific patterns of their activation or repression during Treg differentiation from peritumors to tumors. In line with data at the level of single genes, SCORPIUS was able to clearly separate Tregs from the two different sites (**Fig. 5C**), and identified domains of activity (**Fig. 5D**), suggesting that different TFs might be involved at different steps of Treg hyperactivation in iCCA. Specifically, loss of activity of FLI1, recently shown to inhibit effector CD8+ T cell differentiation in murine models of chronic infection and cancer (Chen et al., 2021), was accompanied by increased activity of several TFs simultaneously, *e.g.*, of IKZF2, SMAD1, VDR, IRF4, FOXP3 and BATF, during transition from peritumors to tumors. Several of these TFs are known to be upregulated, or to play a direct role in the enhanced immunosuppression of Tregs in solid tumors. We also revealed increased activity related to enhancer of zeste homolog 2 (EZH2), a histone H_3_K_27_ methyltransferase elevated in tumor-infiltrating Tregs and whose pharmacological inhibition results in proinflammatory Treg reprogramming and enhanced antitumor immunity (Goswami et al., 2018; Wang et al., 2018). A second group of TFs, including MEOX1, TP73 (encoding p73), SOX9 and FOXA1, among others, was activated transiently in Tregs in iCCA, and was later followed by enhanced ZMIZ1 and MYB activities, the latter reported to regulate effector Treg differentiation in murine peripheral organs (Dias et al., 2017) (**Fig. 5D**). Collectively, our analysis identified novel TF activities possibly related to hyperactivated Treg differentiation and enhanced immune suppression in iCCA.

### MEOX1 transcriptionally and epigenetically reprograms circulating Tregs to a tumorinfiltrating phenotype

We next focused our investigation on one of the top hits from SCENIC analysis of intratumoral Tregs, *i.e.*, MEOX1, whose role in the immune system in unknown. MEOX1 encodes a mesodermal TF that plays a key role in somitogenesis and sclerotome development and whose mutation in humans results in the incomplete development of bones in the neck (also known as Klippel-Feil syndrome) (Mohamed et al., 2013; Skuntz et al., 2009). *MEOX1* mRNA levels strongly correlated with those of *FOXP3* in the cholangiocarcinoma dataset from the cancer genome atlas (TCGA; **Fig. S4A**), suggesting a relationship with Tregs in tumors. In our scRNA-seq dataset, MEOX1 expression could be detected only in tumor-infiltrating Tregs, and not in other infiltrating CD4+ or CD8+ T cells (**Fig. S4B**). Similarly, Tregs overexpressed significantly more MEOX1 than CD4+ Tconv from lung, ovarian and breast tumors (**Fig. S4C**). A similar trend could be found in colorectal tumors, although not reaching statistical significance (**Fig. S4C**). By analysing the MEOX1 promoter, we found binding motifs of FOXP3, IRF4, FOXO1, EZH2 and IKZF2 TFs, among others (**Fig. 6A**). We also found that, in iCCA, the predicted activity of these TFs was co-regulated with that of MEOX1 in a subset of tumor-infiltrating Tregs (**Fig. 6B**), collectively suggesting a role in the regulation of MEOX1 expression. To investigate the functional importance of MEOX1 in specifying the molecular characteristics of tumor-infiltrating Tregs, we isolated peripheral blood CD4+ Tregs from healthy donors, activated them for 24 hours with αCD3/CD28 + IL-2 and transduced them with a lentivirus capable to overexpress the full-length MEOX1 cDNA, or with a mock lentivirus control. Transduced cells, identified by the green fluorescent protein (GFP) reporter, were further purified as CD127– CD25+ by fluorescence-activated cell sorting (FACS) and analysed at the transcriptomic and chromatin accessibility level by bulk RNA-seq and assay for transposase-accessible chromatin using sequencing (ATAC-seq), respectively (**Fig. 6C**). At the chromatin level, we found that genes previously shown to be overexpressed by Tregs in tumors and associated with effector differentiation (*TNFRSF9*, *IL1RN*) (Alvisi et al., 2020) and with disease progression in multiple cancers (*LAYN*) (De Simone et al., 2016), or responsible for IL-10 production by Tregs (*PRDM1*) (Cretney et al., 2011), among others (**Table S7**), were more accessible at multiple genomic sites in MEOX1-transduced vs. mock-transduced Tregs (**Fig. 6D**). Computational analysis of differentially accessible regions in the whole ATAC-seq dataset further identified differentially accessible TF-binding motifs (TFBMs). Motifs attributable to AP-1 activity (FOS, JUNB and the combined FOS:JUNB motif), to AP-1 transcriptional partners (BATF, IRF4 and the combined BATF:JUN motifs, mechanistically linked to hyperactivated Tregs in tumors) (Alvisi et al., 2020; Grant et al., 2020) and the SMAD2:SMAD3 combined motif, reflecting increased accessibility of genes possibly controlled by TGF-β signaling, were enriched in MEOX1-transduced Tregs, whereas motifs attributable to TWIST2, MXI1, KLF9 and TP53 were enriched in mock-transduced Tregs (**Fig. 6E**). In agreement with ATAC-seq results, overexpression of MEOX1 resulted in the induction of *TNFRSF9*, *IL1RN*, *LAYN*, as well as of additional effector or iCCA Treg-related genes such as *CD70*, *MAGEH1* and *ICOS* (**Fig. 6F** and **Fig. S3D**) (Alvisi et al., 2020; De Simone et al., 2016), and of *NRP1* (encoding Neuropilin-1), *IL10* and *CTLA4* (**Fig. 6F**), all involved in Treg-mediated immunosuppression. Gene set enrichment analysis further showed that MEOX1-overexpressing Tregs were strongly enriched in specific transcriptomic signatures of Treg vs. CD4+ Tconv, of iCCA-infiltrating vs. peritumor-infiltrating Tregs, or of ICOS+ CCR8+ vs. ICOS– CCR8– Tregs recently described by our group as highly immunosuppressive in non-small cell lung cancer (NSCLC), melanoma and hepatocellular carcinoma (**Fig. 6G**) (Alvisi et al., 2020). Collectively, these data support the conclusion that TF MEOX1 promotes the acquisition of a tumor-infiltrating Treg phenotype by reprogramming the transcriptional and epigenetic landscape.

**Figure 6.**
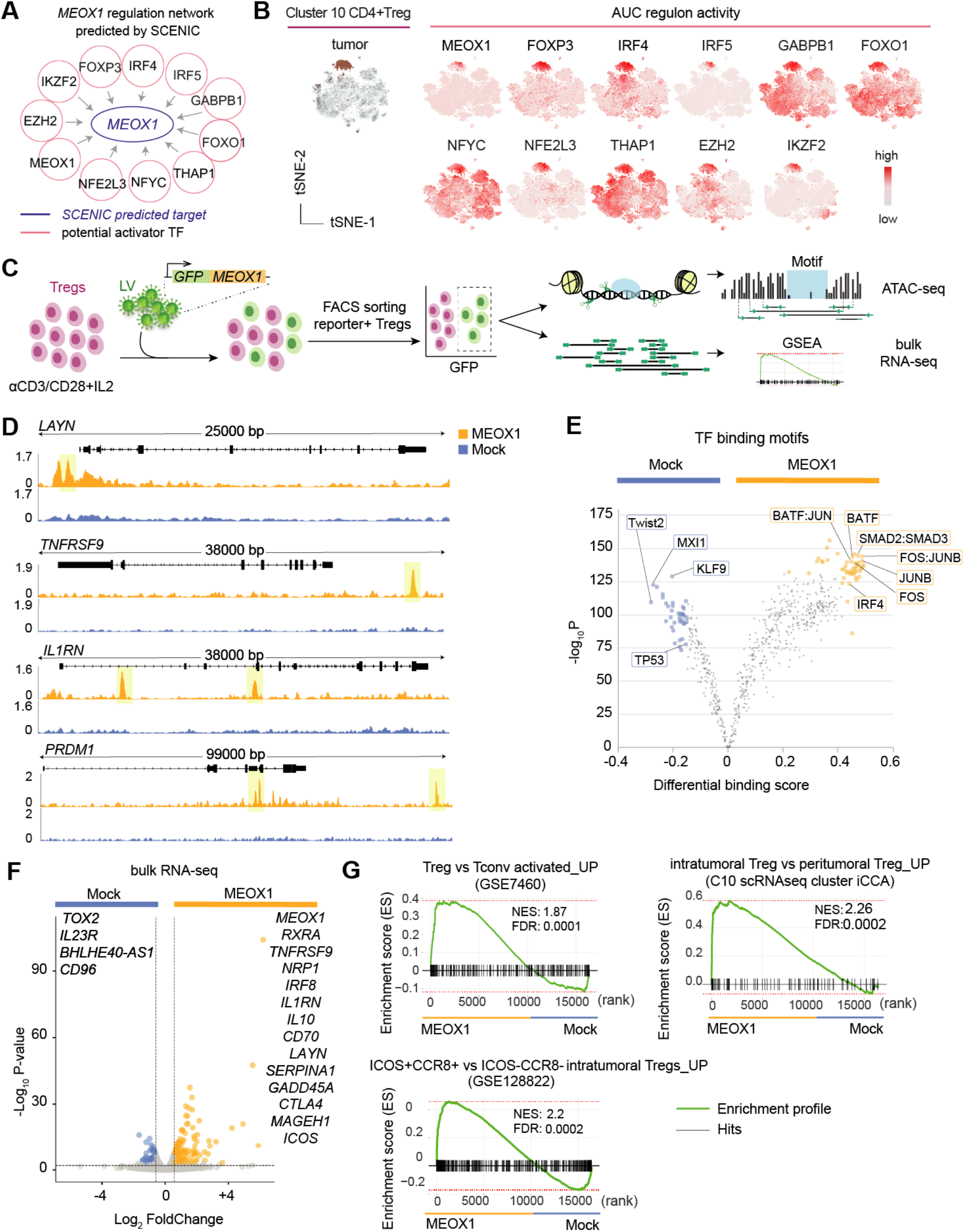
MEOX1 transcriptionally and epigenetically reprograms Tregs to a tumor-infiltrating phenotype. **A.** MEOX1 regulation network predicted by SCENIC based on DNA-motif analysis (RcisTarget database). **B**. t-SNE map of all T cells obtained as in Fig. 5A and depicting Treg cells from tumors (scRNAseq cluster 10) and specific TF activities (predicted activators of MEOX1), as indicated by Area Under the recovery Curve (AUC) values. **C.** Schematic view of the MEOX1 overexpression (OE) approach by lentiviral vector (LV) in Tregs isolated from the peripheral blood of healthy donors (n=3). ATAC-seq: Assay for Transposase-Accessible Chromatin using sequencing; GSEA: Gene Set Enrichment Analysis. **D.** Representative accessible genomic regions in ATAC-seq data from Treg cells transduced as in C. Significant differentially accessible regions are highlighted in light yellow. **E.** Transcription factor binding motif (TFBM) enrichment analysis by TOBIAS. Colored dots represent single significant motifs. TFs of interest are indicated. **F.** Volcano plot of differentially expressed genes in bulk RNA-seq from MEOX1- and mock-transduced Tregs (q-value< 0.05, |FC|> 1.5, n=3). Genes of interest are indicated. **G.** GSEA of different gene sets in bulk GSEA of different gene sets in bulk RNA-seq data obtained as in C.

## Discussion

We provide a comprehensive characterization of T-cell and myeloid subsets that are present in patients with iCCA by high-dimensional single cell technologies and report the extensive infiltration of CD4+ Tregs with a highly immunosuppressive phenotype, that is accompanied by the loss of CD8+ CTLs compared to the peritumoral tissue. CD8+ T cells expressing CD39, a marker recently linked to the identification of tumor-specific CD8+ T cells in the tumor microenvironment, represented only a minor fraction of CD8+ T cells in iCCA, which appears to be much lower than that reported for highly immunogenic tumors, such as NSCLC, melanoma and MSI colon cancer (Duhen et al., 2018). This observation reinforces previous evidence that, overall, iCCA is poorly immunogenic, thereby contributing to explain, among other factors, its low response to checkpoint blockade with anti-PD1 (Kim et al., 2020; Piha-Paul et al., 2020). By contrast, changes at the level of the myeloid compartment were less evident, and were mainly ascribed to the increased infiltration of cDC2, rather than to changes in overall gene expression, compared to other tissue sites. Importantly, absence of these cDC2 identified patients of our cohort with faster disease progression when also Tregs were high, in agreement with a recent report in head and neck squamous cell carcinoma (Binnewies et al., 2019), thus suggesting that a common axis might regulate anti-tumor immunity in different cancer types.

scRNA-seq further informed on the characteristics of the different immune cell types and enabled the identification of a major suppressive hub orchestrated by Tregs in the iCCA tumor microenvironment. Compared to the adjacent peritumoral tissues, iCCA-infiltrating Tregs showed features of the hyperactivated effector state previously described in several solid tumors, including melanoma, NSCLC, hepatocellular carcinoma, breast and colon cancer (Plitas and Rudensky, 2020). Although differences that are yet to be identified might be present according to the specific tumor type, it seems evident that intratumoral Tregs share a core transcriptional and functional program, characterized by the increased expression of inhibitory molecules, chronic immune activation and enhanced suppressive capacity compared to those from the circulation or the peritumoral area. Tregs are predicted to engage multiple inhibitory pathways on T and myeloid cells in the tumor microenvironment, while receiving signals that in turn may support their hyperactivated phenotype, thereby offering novel, more specific targets for cancer immunotherapy. In agreement with the Treg-mediated inhibition, in several iCCA-infiltrating T-cell subsets we observed reduced activity of TFs promoting the cytotoxic program and effector function of T cells, including RUNX3, EOMES and TBX21 activities. We still do not know what is the relationship between enforced surface interactions and altered downstream gene expression/TF activities in T cells in iCCA. We anticipate that the future development of computational tools capable to integrate multiple layers of information, including at the level of single cells, will identify with precision those regulatory circuits that should be targeted to interfere with specific dysfunctional programs. Among infiltrating T cells, Tregs had the most diverse molecular program compared to those present in the adjacent peritumoral tissue, corroborating the hypothesis that these cells play a central role in orchestrating immunosuppression iCCA. We identified several TFs whose enhanced activity has been previously demonstrated to regulate immune activation of Tregs in solid tumors. These included IRF4, BATF and EZH2, which play an essential role in suppression of anti-tumor immunity (Alvisi et al., 2020; Goswami et al., 2018; Wang et al., 2018), in addition to TFs whose functional role in Treg biology remained to be established. Among these, we found that overexpression of MEOX1 was sufficient to epigenetically and transcriptionally reprogram circulating Tregs to a tumor-infiltrating phenotype. The mechanisms by which MEOX1 is induced and operates in hyperactivated Tregs remains to be established. MEOX1 is triggered by TGF-β signaling in non-immune cells such as mesenchymal progenitors and cardiac fibroblasts (Alexanian et al., 2021; Dong et al., 2018), and mediates cell differentiation by direct binding to DNA. Although these aspects were not tested directly on intratumoral Tregs, pseudotemporal ordering of TF activities and molecular analyses inspire a model according to which IRF4 and BATF precede MEOX1 activity, which in turn sustains hyperactivated Treg gene expression and, possibly, immunosuppression by favoring chromatin accessibility at AP-1/IRF4/BATF consensus sites.

In conclusion, we provide a comprehensive characterization of the iCCA T cell and myeloid infiltrate, and show that Treg abundantly infiltrate the tumor microenvironment while CTLs and CD39+, putative tumor-specific CD8+ T cells expressing PD-1 are rare. Future immunotherapeutic strategies must thus aim to turn iCCA from cold to hot, so to favor T-cell infiltration. Interfering with the hyperactivated Treg program, such as that regulated by MEOX1, which is shared by several solid tumors, seems an attractive approach to consider in this regard.

## MATERIALS AND METHODS

### Study design

The use of human samples was approved by the Humanitas Research Hospital Institutional Review Board (approval no. 146/20 for patients’ samples and no. 28/01/2016 for buffy coats from healthy donors). All patients were enrolled at Humanitas Research Hospital and provided informed consent for tissue biobanking. All patients had pathology proven iCCA, and were without chemotherapy treatment prior to resection. Hepatitis B or C positive patients were excluded. Samples (one tumoral and one peritumoral) were selected from areas without macroscopic evidence of necrosis or hemorrhage. For morphological analysis, sections were cut (2μm thick), stained with hematoxylin and eosin and evaluated by an expert liver pathologist. The histological features that were evaluated included tumor grade, desmoplasia, resection margin, steatosis, perineural, and linfo-vascular invasion, and lymph node metastasis. Resected iCCA patients were followed up every 3 months, as per protocol in our Center, or until death, and major events were recorded. Characteristics of the patients enrolled in the study are listed in **Supplementary Table 1**.

### Sample collection and processing

PBMCs were isolated from buffy coats from iCCA and healthy donors via density-gradient separation and were cryopreserved in FBS supplemented with 10% DMSO until use. Blood samples from patients were collected in vacutainer EDTA Tubes (BD). Tissues were collected in complete R10 medium: RPMI-1640 medium with 10% FBS (Sigma-Aldrich), 1% penicillin-streptomycin and 1% Ultra-glutamine (both from Lonza) RPMI 1640 media: 10% FBS (Sigma-Aldrich), 1% penicillin-streptomycin (Lonza) and 1% Ultra-glutamine (Lonza). Human cell isolation from healthy liver was performed by collagenase perfusion adapting a traditional two-step technique (Seglen, 1976). Briefly, non-tumoral tissue displaying the normal vascular architecture, was perfused in order to collect the circulating intrahepatic blood (perfusate). Then, the liver sample was digested with collagenase, obtaining a single-cell suspension. Parenchymal cells were eliminated from the sample through appropriate centrifugations and the supernatant was counted and stored. Cells were pelleted, counted, and frozen according to the slow freezing procedure in standard cryo-vials. Further disaggregation of iCCA tissue into a single-cell solution for sequencing was completed using the MACS tumor dissociation kit (Miltenyi Biotec) with the standard tough tumor protocol. Briefly, the MACS tumor dissociation kit enzyme mix (300μl) was added to each sample. Next, samples were put into the gentleMACS Dissociator and ran through the tough tumor program. The cell suspension was then applied to a 70um cell strainer. Cells were pelleted, counted and frozen according to the slow freezing procedure in standard cryo-vials (1 ml cell suspension in the cryopreservation medium as above.

### Polychromatic flow cytometry

Frozen samples were thawed in R10 additioned with 20 μg/mL DNase I from bovine pancreas (Sigma-Aldrich). After washing with PBS (Sigma-Aldrich), the cells were stained with the combination of mAbs listed in **Supplementary Table 2**. Flow cytometry procedure for (i) selection of antigen-fluorochrome combinations, (ii) reagent titration and validation, (iii) limiting spreading error (SE) have been previously described (Brummelman et al., 2019). Flow cytometry data were analysed on a FACS Symphony A5 cytometer (BD Biosciences) equipped with 5 lasers (UV, 350 nm; violet, 405 nm; blue, 488 nm; yellow/green, 561 nm; red, 640 nm; all tuned at 100 mW, except for UV, tuned at 60 mW). Compensation was performed by FlowJo by using BD Compbeads inclubated with single antibodies, as previously described (Lugli et al., 2017).

### Computational analysis of flow cytometry data

Data were processed as previously described (Brummelman et al., 2019). Briefly, Flow Cytometry Standard (FCS) 3.0 files were analysed by standard gating in FlowJo version 9 to remove dead cells and spurious events, and identify CD45+CD3+ T cells or CD45+CD3-CD66b-myeloid cells. 3,000 CD45+CD3+ T cells and 2,000 CD45+CD3-CD66b-myeloid cells per sample were subsequently imported in FlowJo (version 10), biexponentially transformed, and exported for further analysis by a custom-made pipeline of PhenoGraph where we modified the Linux-community and the core.py script of PhenoGraph package to fix the seed to “123456” (run in Python version 3.7.3). Tumoral, peritumoral, perfused blood from tissue and PBMC samples were labelled with a unique computational barcode for further identification and converted in comma separated (CSV) files and concatenated in a single matrix by using the merge function of pandas package. The K value, indicating the number of nearest neighbours identified in the first iteration of the algorithm, was set at 100 and 200 for CD3^+^ and for CD3-CD66b-cell clustering, respectively. The data were then reorganized and saved as new CSV files, one for each cluster, that were further analysed in FlowJo to determine the frequency of positive cells for each marker and the corresponding median fluorescent intensity (MFI). Subsequent metaclustering of these values was performed using the gplots R package. Uniform Manifold Approximation and Projection (UMAP) was obtained by UMAP Python package.

### scRNA-seq

Frozen tumoral and peritumoral single-cell suspension were thawed, washed in PBS, stained with Live/dead Aqua Fluorescent Reactive Dye (Life Technologies) and mouse anti-human CD45 antibody (30-F11, BD Biosciences) at 4°C for 20 minutes and sorted on a FACS Aria III (BD Biosciences) with a 100μm nozzle. FACS-purified CD45^+^/CD45^−^ cells were resuspended in 1ml PBS plus 0.04% BSA and washed two times by centrifugation at 450xg for 7min. After the second wash, cells were resuspended in 30 μl and counted with an automatic cell counter (Countess II, Thermo Fisher) to get a precise estimation of total number of cells recovered and of their concentration. Afterwards, CD45^+^ cells of each sample were loaded into one channel of the Single Cell Chip A using the Single Cell 3’ reagent kit v2 single cell reagent kit (10X Genomics) for Gel bead Emulsion generation into the Chromium system. Following capture and lysis, cDNA was synthesized and amplified for 14 cycles following the manufacturer’s protocol (10X Genomics). 50 ng of the amplified cDNA were then used for each sample to construct Illumina sequencing libraries. Sequencing was performed on the NextSeq550 Illumina sequencing platform following 10x Genomics instruction for reads generation. A sequencing depth of at least ~ 30,000 reads/cell was obtained for each sample.

### Pre-processing of scRNA-seq data

Raw sequencing data was processed and aligned to the GRCh38 human reference genome with CellRanger (10X Genomics) v3.0.1 (Zheng et al., 2017). Resulting filtered matrices of molecular counts (count matrices) were used as input for pre-processing with Seurat package v3.0.3 (Butler et al., 2018) running under R v3.6.1 and Bioconductor v3.9 on a Debian GNU/Linux 9 operating system. First, ribosomial genes were removed from the count matrix. Then, in order to avoid dying/damaged cells or doublets, quality control was performed and cells having < 200 or > 3000 expressed genes or >20% mitochondrial counts were filtered out. Data normalization and log-transformation was performed by applying the NormalizeData Seurat function (method = LogNormalize). Cell cycle scores were then evaluated by calling the CellCycleScoring Seurat function using a list of cell cycle markers genes (Tirosh et al., 2016).

### Dataset Integration

In order to integrate sample datasets together, the “SCT” normalization (Hafemeister and Satija, 2019) was performed separately for each sample by running the SCTransform Seurat function where technical and cell-cycle effects were regressed out (do.correct.umi = true; vars.to.regress = percent.mt, S.Score, G2M.Score). The Seurat functions PrepSCTIntegration, FindIntegrationAnchors and IntegrateData were then applyed on the list of Seurat objects using 3,000 genes in the anchor finding process. A final integrated dataset consisting of 31,745 cells (14,824 from peritumoral and 16,921 from tumoral samples) was obtained. A principal component analysis (PCA) was performed on the top variable features by calling the RunPCA Seurat function and the first 20 principal components (PCs) were selected for downstream analysis. Imputation of missing count values was performed by applying the Adaptively-thresholded Low Rank Approximation (ALRA) method (Linderman et al., 2018) by calling the RunALRA Seurat function with default parameters.

### Clustering and cell type identification

To identify cell clusters, the graph-based clustering approach implemented in the FindClusters Seurat function was used with a resolution value ranging from 0 to 1 by steps of 0.1. Clustering stability was assessed using clustree R package v.0.4.0 (Zappia and Oshlack, 2018). Marker genes were obtained for each cluster by running the FindAllMarkers Seurat function which performs a Wilcoxon Rank Sum test (adj.p-value < 0.05). The SingleR v1.0.6 (Aran et al., 2019) and AUCell v1.8.0 (Aibar et al., 2017) packages were used as an aid to manual clusters cell types annotation.

### Reclustering of T cells

Reclustering of T cells was performed by repeating the analysis used for the whole dataset on the subsets of cells enriched for the “Main Immune Cell Expression Data”’ CD8 and CD4 T cells reference signature after ALRA imputation (5,037 cells from peritumoral and 7,607 cells from tumoral samples). A clustering resolution of 0.8 was chosen, obtaining a total of 11 clusters.

### Reclustering of myeloid cells

Reclustering of myeloid cells was performed by repeating the analysis used for the whole dataset, apart from lowering k.filter to 100 in FindIntegrationAnchors Seurat function, on the subsets of cells enriched for the “Main Immune Cell Expression Data’” myeloid reference signature after ALRA imputation (2,384 cells from peritumoral and 2,201 cells from tumoral samples). A clustering resolution of 0.5 was chosen, obtaining a total of 8 clusters.

### UMAP plots

UMAP was obtained by running the RunUMAP Seurat function with default parameters for the whole dataset and with min.dist = 0.01 and n.neighbors = 20 for reclusterings of T cells and Myeloid cells.

### PCA of samples by gene expression values

Averaged gene expression values were obtained by running the AverageExpression function on the RNA assay of the Seurat object with data being normalized, centered and scaled by using the NormalizeData (method = LogNormalize) and ScaleData Seurat functions. PCA was performed with FactoMineR package v2.3. The first two principal components were plotted using factoextra package v1.0.6.

### Heatmap of cluster marker genes

Cluster marker genes were obtained by running the FindAllMarkers Seurat function after imputation of missing count values with ALRA algorithm as previously described. For each cluster, genes having an adjusted p-value < 0.05 and absolute log2 of fold-change (FC) > 0.5 and detectable expression in > 50% of the cells in that cluster were selected and sorted in descending order by FC values (**Supplementary table 3**). After data centering and scaling with ScaleData, the heatmap was generated using the DoHeatmap Seurat function with default parameters.

### Differentially expressed genes (DEGs) from scRNA-seq

For each cluster, DEGs were obtained by comparing cells from peritumoral samples with cells from tumoral samples using the FindMarkers Seurat function (min.pct = 0.1, logfc.threshold = 0), which performs a Wilcoxon Rank Sum test. Genes having an adjusted p-value < 0.01 and an absolute FC > 1.5 were considered as statistically significant. Volcano plot was generated with EnhancedVolcano package v1.4.0 (https://github.com/kevinblighe/EnhancedVolcano).

### Integration of publicly available scRNA-seq datasets

To generate **Fig. S4C**, we processed scRNA-seq data on lung (n=3), breast (n =14), ovarian (n =5) and colorectal (n =7) tumors by Qian et al. (https://lambrechtslab.sites.vib.be/en/pan-cancer-blueprint-tumour-microenvironment-0) (Qian et al., 2020). We filtered out cells labelled as “Normal” and the 48,231 cells containing a mixture of different T-cell phenotypes were normalized (target_sum=1e4) using ‘‘scanpy.pp.normalize_total’’ function from SCANPY package (Wolf et al., 2018). CPM counts were square-root-transformed (“scanpy.pp.sqrt”) followed by dimensionality reduction by PCA on top most variable genes, computed using “scanpy.pp.highly_variable_genes” function with the following parameters: min_mean=0.0125, max_mean=3, min_disp=0.5. De novo clustering was performed using “scanpy.tl.leiden” function and marker genes for each significant cluster were found using the function “scanpy.tl.rank_genes_groups” (method= Wilcoxon). CD4+ T cell clusters were isolated and imputed with the MAGIC algorithm (k, number of nearest neighbors = 5) (van Dijk et al., 2018). Tregs were defined as those cells belonging to the cluster with the highest expression of Treg-related genes. The remaining CD4+ T cells were combined and labelled as Tconv. The box plots showing the median expression of imputed transformed data were generated using seaborn package (version 0.11.1) and the statistical significance were assessed with Mann-Whitney-Wilcoxon test function of the statsmodels package (version= 0.12.2).

### Cell-cell interaction analysis

Cellular interactions between Treg and myeloid/T cell populations were analysed with CellphoneDB tool (version 2.0.0) with default parameters (Vento-Tormo et al., 2018). Normalized gene expression matrices together with their cluster annotations derived from both myeloid and T cell reclustering were used as input (‘data’ slot from Seurat after ALRA imputation step). The enriched ligand-receptor interactions between two cell subsets were calculated based on permutation test. Fig. 4 lists L:R pairs with a p-value ≤0.05, manually curated from all interactions listed in **Supplementary Table 4**.

### Gene regulatory network analysis

The analysis of regulon activity were performed by SCENIC (version 1.1.2) (Aibar et al., 2017) starting from ALRA imputed data. The genes with at least 3 UMIs in at least 10% of the cells and detected in at least 10% of samples were selected as the input genes. The expression matrix was loaded into GENIE3 (Huynh-Thu et al., 2010) and the co-expressed genes to each TF was constructed. The TF co-expression modules were then analysed by RcisTarget Bioconductor package. The Normalized Enrichment Score (NES) of the transcription factor binding motifs (TFBS) was calculated and NES > 3.0 were considered as significantly enriched. The filtered potential targets by RcisTarget mouse hg19 database (hg19-500bp-upstream-7species.mc9nr.feather; hg19-tss-centered-10kb-7species.mc9nr.feather) from the co-expression module were used to build the regulons. The regulon activity was analysed by AUCell (Aibar et al., 2017) and the active regulons were determined by AUCell default threshold. The active regulons were then mapped to all cells by *t*-SNE in Fig. 5A. Binary regulon activity active in at least 50% of cells were selected. Binary regulon activity clustering analysis was performed for Figure 5B. The complete list of significant regulons is listed in **Supplementary Table 5**.

### Trajectory analysis

Trajectory analysis was performed with SCORPIUS R package v1.0.7 (Saelens et al., 2019) on Regulon Activity Scores obtained by running SCENIC R package as described in the previous paragraph. Dimensionality reduction was obtained by applying the reduce_dimensionality function (dist = “spearman”, ndim = 3) on the subset of cells belonging to C10 of T cells. Linear trajectory was inferred by using the infer_trajectory function with default parameters. The importance of each gene with respect to the trajectory was calculated by running the gene_importances function (num_permutations = 10, ntree = 10000, ntree_perm = 1000). Regulons having a false discovery rate (FDR)-adjusted p-values < 0.05 were considered significant and are listed in **Supplementary Table 6**.

### Treg cell transduction and culture condition

The use of recombinant plasmids was approved by the Italian Ministry of Health authorization number MI/IC/Op2/17/024. Treg cells were enriched from buffy coats of healty donors using an EasySep Human CD4+ CD127^low^ CD25+ Regulatory T cell Isolation kit. Purity was confirmed to be >90% by flow cytometry. Purified Treg cells were stimulated with Dynabeads human T-ACT CD3/CD28 at a 1:2 bead:cell ratio, in 96 U-bottomed well plates and in the presence of IL-2 (50 ng/mL; Peprotech). 24h and 48h after stimulation, cells were transduced with lentiviral particles harboring a custom human ORF for MEOX1 (Sigma-Aldrich, Mission TRC3) or empty backbone-vectors (mock), both expressing GFP as a reporter (Sigma-Aldrich). Cells were cultured for 5 additional days and then stained with fluorochrome-conjugated monoclonal antibodies (listed in **Supplementary Table 2**). GFP+ cells, pre-gated as CD3+CD8-CD4+Aqua-CD25+CD127-were isolated with a FACSAria cell sorter (BD Biosciences).

### ATAC-seq

Libraries were prepared using a protocol adapted from Buenrostro et al. (Buenrostro et al., 2015), as previously described (Galletti et al., 2020). 10,000 FACS-purified GFP^+^ Treg cells were washed in PBS without Ca^2+^ and Mg^2+^ and resuspended in 25μl lysis buffer (10 mM Tris-HCl pH 7.4, 10 mM MgCl2, 0.1% IPEGAL CA-630). Nuclei were pelleted by centrifugation for 10 min at 500g and resuspended in a final reaction volume of 25μl comprising 0.2μl of Tn5 transposase (made inhouse), 5μl of 5× transposase buffer (50 mM Tris-HCl pH 8.4, 25 mM MgCl2) and 19.8μl of ultrapure water (Milli-Q). The reaction was incubated with mixing at 300 r.p.m. for 30 min at 37 °C, supplemented with 5 μl clean-up buffer (900 mM NaCl, 30 mM EDTA), 2.5μl of 20% SDS, 0.35μl of ultrapure water (Milli-Q) and 2.15μl of proteinase K (18.6 μg μl–1, ThermoFisher Scientific), and incubated for a further 30 min at 40 °C. Tagmented DNA was isolated using 2× SPRI Beads (Beckman Coulter) and amplified via PCR. Fragments smaller than 600 bp were isolated via negative size selection using 0.65× SPRI Beads (Beckman Coulter) and purified using 1.8× SPRI Beads (Beckman Coulter). Quality control was performed using a 4200 TapeStation System (Agilent) in conjunction with a Qubit 2.0 Fluorometer (Thermo Fisher Scientific). Libraries were then multiplexed in an equimolar pool and sequenced using a NextSeq 500/550 Platform (Illumina). At least 40 million single-end 75-bp reads were generated per sample.

### ATAC-seq data analysis

After demultiplexing with bcl2fastq v2.20 (Illumina, Inc.), quality control checks on raw sequencing data were performed with FastQC v0.11.8 (https://www.bioinformatics.babraham.ac.uk/projects/fastqc/). Adapters removal and dynamic trimming of low-quality bases were performed using Trimmomatic v0.39 (Bolger et al., 2014) with parameters “ILLUMINACLIP:TruSeq2-SE.fa:2:30:10, LEADING:3, SLIDINGWINDOW:4:15, MINLEN:36”. Single-end reads were mapped to the UCSC hg38 reference human genome using Bowtie2 v2.4.1 (Langmead and Salzberg, 2012) with parameters “--end-to-end -D 20 -R 3 -N 1 -L 20 -i S,1,0.50 --no-unal”. After alignment, several post-processing has been carried out. First, PCR duplicates, reads aligned to chrM and/or encode blacklisted regions v2 (Amemiya et al., 2019) were filtered out using BEDTools v2.30 (Quinlan and Hall, 2010). Second, only uniquely mapped reads were retained using SAMtools v1.9 (Li et al., 2009). Finally, reads aligning to the forward strand were offset by +4 bp, and reads aligning to the reverse strand were offset −5 bp. Non-shifted bam files were also kept for the differentially bound motif analysis. Peak calling was performed using MACS2 v2.2.6 (Zhang et al., 2008) with parameters “--nomodel --extsize 200 --shift −100 --gsize 2.9e9 --tsize 71 --qvalue 0.05 -bdg --SPMR --keep-dup all”. In order to visualize the raw profiles with the Integrative Genomics Viewer (IGV) v2.9 (Robinson et al., 2011), RPM-normalized BedGraph files generated by MACS2 were converted to BigWig files v2.8 from UCSC tools (Kent et al., 2010). In order to obtain a set of unique genomic regions for the two conditions, MACS2 called peaks were processed with BEDTools v2.30. Each region was extended by 50bp at both ends and theirs overlapping mapped reads were counted with Rsubread v2.4.2 (Liao et al., 2019) R package. The obtained count matrix was imported and processed by DeSeq2 v1.30.1 (Love et al., 2014) R package, and differential expression analysis was performed with a paired design. Differentially significant regions between the two conditions were selected by choosing a p-value cut-off of 0.05 (**Supplementary Table 7**).

### Volcano plot of differentially bound motifs

Differentially bound motifs were obtained by applying the TOBIAS v0.12.10 pipeline (Bentsen et al., 2020) to unshifted bam files obtained as described previously. Briefly, peak-calling was performed per replicate using MACS2 v2.2.6 with parameters “--nomodel --extsize 200 --shift −100 --gsize 2.9e9 --tsize 71 --broad --qvalue 0.01 --keep-dup ‘all’” and the list of genomic regions that represents peaks merged across all conditions was obtained using BEDTools v2.30. Then, single replicates bam files were merged to condition bam files with SAMtools v1.9. The TOBIAS core analysis was then performed. First, to correct the Tn5 transposase insertion bias and to calculate a continuous footprinting score across regions, the ATACorrect and FootprintScores tools were runned for each condition with default parameters. Second, to make predictions on specific TF binding sites and to obtain the differential binding between conditions, the BINDetect tool was called with default parameters and by using the JASPAR 2020 core vertebrates non-redundant set of TF binding profiles (Fornes et al., 2020) (**Supplementary Table 8**).

### Bulk RNA sequencing (RNA-seq)

RNA isolation of the FACS-purified GFP^+^ Treg cells was performed by the Quick-RNA Microprep kit following the manufacturer’s protocol (Zymo Research). RNA quality control was performed with the Agilent 4200 Tape Station system and only RNAs having a RIN >8 were used for library preparation. Libraries for mRNA sequencing were prepared starting from 10 ng tot RNA for each sample by using the SMART-Seq v4 Ultra Low Input RNA Kit (Clontech-Takara). All samples were sequenced on an Illumina NextSeq 550 at an average of 17,5 million 75-bp single-end reads.

### Bulk RNA-seq data analysis

After demultiplexing with bcl2fastq v2.20 (Illumina, Inc.), quality control checks on raw sequencing data were performed with FastQC v0.11.8 (https://www.bioinformatics.babraham.ac.uk/projects/fastqc/)). Adapters removal and dynamic trimming of low-quality bases were performed using Trimmomatic v0.39 with parameters “ILLUMINACLIP:TruSeq2-SE.fa:2:30:10, LEADING:3, MAXINFO:50:0.7, MINLEN:40”. Single-end reads were aligned to the UCSC hg38 human genome using STAR v2.7.0f (Dobin and Gingeras, 2015) with parameters “--outSAMtype BAM SortedByCoordinate, --outFilterMultimapNmax 20, --outWigType wiggle, --outWigNorm RPM”. Wiggle files were converted to BigWig format using wigToBigWig v2.8 from UCSC tools, Uniquely mapped reads having MAPQ > 30 were selected for downstream analysis using SAMtools v1.9. Quality control checks of aligments were carried out using Qualimap v2.2.2a (Okonechnikov et al., 2016). Transcripts quantification was performed by running the featureCounts function of the Rsubread package v1.34 with parameters “minOverlap=5, isPairedEnd=FALSE, strandSpecific=0”. Raw count values were then loaded and processed within edgeR R package v3.28.1 running under R v3.6.3 and Bioconductor v3.9. Non-expressed genes were filtered out by keeping only genes with read counts greater than 1 Count Per Million (CPM) in at least one sample. Then, data was normalized by applying the Trimmed Mean of M-values (TMM) normalization. Differential expression analysis was performed using the GLM approach using a paired design, and significant DEGs were obtained by calling the topTable function by choosing a minimum absolute FC of 1.5 and FDR q-value < 0.05 (**Supplementary Table 9**).

### Volcano plot of DEGs from RNA-seq

Volcano plot was generated with EnhancedVolcano package v1.4.0 (https://github.com/kevinblighe/EnhancedVolcano). Genes having an adjusted p-value < 0.05 and an absolute FC > 1.5 were considered as statistically significant.

### Overrepresentation analysis

Gene set enrichment analysis (GSEA) was obtained by running the fgsea R package v1.12.0 (Korotkevich et. al, 2019), which perform a pre-ranked GSEA. The function fgsea was applied on the ranked list of all genes with default parameters and “maxSize=1000, nperm=10000”. Ranking score was determined by a combination of fold-change and F-statistic. Gene sets evaluated included: (*i*) Treg vs Tconv activated_UP (GSE7460) gene set (Hill et al., 2007); (*ii*) ICOS+CCR8+ vs ICOS-CCR8-intratumoral Treg_UP (GSE128822) gene set (Alvisi et al., 2020); (*iii*) intratumoral Treg vs peritumoral Treg_UP (C10 of CD4^+^ Tregs from scRNA-seq).

## Data availability

Raw data sets are available in the Gene Expression Omnibus (GEO) database (http://www.ncbi.nlm.nih.gov/geo) under accession number GSE171900, which comprises scRNA-seq data (GSE171899) and RNA-seq data (GSE171895). ATAC-seq data is provided upon request.

## Code availability

Scripts used to analyse the flow cytometry single-cell data are available at https://github.com/luglilab/Cytophenograph. All other codes are available on request.

## Acknowledgements

The authors wish to thank all the patients who participated to this study and G. Natoli and S. Polletti (European Institute of Oncology, Milan) for assistance with the ATAC-seq protocol. This work was funded by the Associazione Italiana per la Ricerca sul Cancro (AIRC IG 2017 – ID 20676 to E.L. and AIRC IG 2019 – ID 23408 to A.L.). G.G. and S.P. were supported by Fellowships from the Fondazione Italiana per la Ricerca sul Cancro-Associazione Italiana per la Ricerca sul Cancro (FIRC-AIRC). K.P. is a recipient of the Fondazione Umberto Veronesi 2020 postdoctoral fellowship. D.D. was supported by Associazione Italiana per la Ricerca sul Cancro AIRC (AIRC Start up 19141) and Minsal (GR-2016-02363531). The purchase of a FACSSymphony A5 was defrayed in part by a grant from the Italian Ministry of Health (Agreement 82/2015).

## REFERENCES

Abou-Alfa, G. K., Macarulla, T., Javle, M. M., Kelley, R. K., Lubner, S. J., Adeva, J., Cleary, J. M., Catenacci, D. V., Borad, M. J., Bridgewater, J., et al. (2020a). Ivosidenib in IDH1-mutant, chemotherapy-refractory cholangiocarcinoma (ClarIDHy): a multicentre, randomised, double-blind, placebo-controlled, phase 3 study. Lancet Oncol 21, 796–807. 10.1016/S1470-2045(20)30157-1

Abou-Alfa, G. K., Sahai, V., Hollebecque, A., Vaccaro, G., Melisi, D., Al-Rajabi, R., Paulson, A. S., Borad, M. J., Gallinson, D., Murphy, A. G., et al. (2020b). Pemigatinib for previously treated, locally advanced or metastatic cholangiocarcinoma: a multicentre, open-label, phase 2 study. Lancet Oncol 21, 671–684. 10.1016/S1470-2045(20)30109-1

Aibar, S., Gonzalez-Blas, C. B., Moerman, T., Huynh-Thu, V. A., Imrichova, H., Hulselmans, G., Rambow, F., Marine, J. C., Geurts, P., Aerts, J., et al. (2017). SCENIC: single-cell regulatory network inference and clustering. Nat Methods 14, 1083–1086. 10.1038/nmeth.4463

Alexanian, M., Przytycki, P. F., Micheletti, R., Padmanabhan, A., Ye, L., Travers, J. G., Gonzalez-Teran, B., Silva, A. C., Duan, Q., Ranade, S. S., et al. (2021). A transcriptional switch governs fibroblast activation in heart disease. Nature. 10.1038/s41586-021-03674-1

Alvisi, G., Brummelman, J., Puccio, S., Mazza, E. M., Tomada, E. P., Losurdo, A., Zanon, V., Peano, C., Colombo, F. S., Scarpa, A., et al. (2020). IRF4 instructs effector Treg differentiation and immune suppression in human cancer. J Clin Invest 130, 3137–3150. 10.1172/JCI130426

Amemiya, H. M., Kundaje, A., and Boyle, A. P. (2019). The ENCODE Blacklist: Identification of Problematic Regions of the Genome. Sci Rep 9, 9354. 10.1038/s41598-019-45839-z

Aran, D., Looney, A. P., Liu, L., Wu, E., Fong, V., Hsu, A., Chak, S., Naikawadi, R. P., Wolters, P. J., Abate, A. R., et al. (2019). Reference-based analysis of lung single-cell sequencing reveals a transitional profibrotic macrophage. Nat Immunol 20, 163–172. 10.1038/s41590-018-0276-y

Banales, J. M., Marin, J. J. G., Lamarca, A., Rodrigues, P. M., Khan, S. A., Roberts, L. R., Cardinale, V., Carpino, G., Andersen, J. B., Braconi, C., et al. (2020). Cholangiocarcinoma 2020: the next horizon in mechanisms and management. Nat Rev Gastroenterol Hepatol 17, 557–588. 10.1038/s41575-020-0310-z

Bentsen, M., Goymann, P., Schultheis, H., Klee, K., Petrova, A., Wiegandt, R., Fust, A., Preussner, J., Kuenne, C., Braun, T., et al. (2020). ATAC-seq footprinting unravels kinetics of transcription factor binding during zygotic genome activation. Nature communications 11, 4267. 10.1038/s41467-020-18035-1

Binnewies, M., Mujal, A. M., Pollack, J. L., Combes, A. J., Hardison, E. A., Barry, K. C., Tsui, J., Ruhland, M. K., Kersten, K., Abushawish, M. A., et al. (2019). Unleashing Type-2 Dendritic Cells to Drive Protective Antitumor CD4(+) T Cell Immunity. Cell 177, 556–571 e516. 10.1016/j.cell.2019.02.005

Binnewies, M., Roberts, E. W., Kersten, K., Chan, V., Fearon, D. F., Merad, M., Coussens, L. M., Gabrilovich, D. I., Ostrand-Rosenberg, S., Hedrick, C. C., et al. (2018). Understanding the tumor immune microenvironment (TIME) for effective therapy. Nat Med 24, 541–550. 10.1038/s41591-018-0014-x

Bolger, A. M., Lohse, M., and Usadel, B. (2014). Trimmomatic: a flexible trimmer for Illumina sequence data. Bioinformatics 30, 2114–2120. 10.1093/bioinformatics/btu170

Borghaei, H., Paz-Ares, L., Horn, L., Spigel, D. R., Steins, M., Ready, N. E., Chow, L. Q., Vokes, E. E., Felip, E., Holgado, E., et al. (2015). Nivolumab versus Docetaxel in Advanced Nonsquamous Non-Small-Cell Lung Cancer. N Engl J Med 373, 1627–1639. 10.1056/NEJMoa1507643

Brummelman, J., Haftmann, C., Nunez, N. G., Alvisi, G., Mazza, E. M. C., Becher, B., and Lugli, E. (2019). Development, application and computational analysis of high-dimensional fluorescent antibody panels for single-cell flow cytometry. Nat Protoc 14, 1946–1969. 10.1038/s41596-019-0166-2

Brummelman, J., Mazza, E. M. C., Alvisi, G., Colombo, F. S., Grilli, A., Mikulak, J., Mavilio, D., Alloisio, M., Ferrari, F., Lopci, E., et al. (2018). High-dimensional single cell analysis identifies stem-like cytotoxic CD8(+) T cells infiltrating human tumors. J Exp Med 215, 2520–2535. 10.1084/jem.20180684

Buenrostro, J. D., Wu, B., Litzenburger, U. M., Ruff, D., Gonzales, M. L., Snyder, M. P., Chang, H. Y., and Greenleaf, W. J. (2015). Single-cell chromatin accessibility reveals principles of regulatory variation. Nature 523, 486–490. 10.1038/nature14590

Butler, A., Hoffman, P., Smibert, P., Papalexi, E., and Satija, R. (2018). Integrating single-cell transcriptomic data across different conditions, technologies, and species. Nat Biotechnol 36, 411–420. 10.1038/nbt.4096

Chen, Z., Arai, E., Khan, O., Zhang, Z., Ngiow, S. F., He, Y., Huang, H., Manne, S., Cao, Z., Baxter, A. E., et al. (2021). In vivo CD8(+) T cell CRISPR screening reveals control by Fli1 in infection and cancer. Cell 184, 1262–1280 e1222. 10.1016/j.cell.2021.02.019

Cretney, E., Xin, A., Shi, W., Minnich, M., Masson, F., Miasari, M., Belz, G. T., Smyth, G. K., Busslinger, M., Nutt, S. L., and Kallies, A. (2011). The transcription factors Blimp-1 and IRF4 jointly control the differentiation and function of effector regulatory T cells. Nat Immunol 12, 304–311. 10.1038/ni.2006

Cruz-Guilloty, F., Pipkin, M. E., Djuretic, I. M., Levanon, D., Lotem, J., Lichtenheld, M. G., Groner, Y., and Rao, A. (2009). Runx3 and T-box proteins cooperate to establish the transcriptional program of effector CTLs. J Exp Med 206, 51–59. 10.1084/jem.20081242

De Simone, M., Arrigoni, A., Rossetti, G., Gruarin, P., Ranzani, V., Politano, C., Bonnal, R. J., Provasi, E., Sarnicola, M. L., Panzeri, I., et al. (2016). Transcriptional Landscape of Human Tissue Lymphocytes Unveils Uniqueness of Tumor-Infiltrating T Regulatory Cells. Immunity 45, 1135–1147. 10.1016/j.immuni.2016.10.021

Dias, S., D’Amico, A., Cretney, E., Liao, Y., Tellier, J., Bruggeman, C., Almeida, F. F., Leahy, J., Belz, G. T., Smyth, G. K., et al. (2017). Effector Regulatory T Cell Differentiation and Immune

Homeostasis Depend on the Transcription Factor Myb. Immunity 46, 78–91. 10.1016/j.immuni.2016.12.017

Dobin, A., and Gingeras, T. R. (2015). Mapping RNA-seq Reads with STAR. Curr Protoc Bioinformatics 51, 11 14 11–11 14 19. 10.1002/0471250953.bi1114s51

Dong, K., Guo, X., Chen, W., Hsu, A. C., Shao, Q., Chen, J. F., and Chen, S. Y. (2018). Mesenchyme homeobox 1 mediates transforming growth factor-beta (TGF-beta)-induced smooth muscle cell differentiation from mouse mesenchymal progenitors. J Biol Chem 293, 8712–8719. 10.1074/jbc.RA118.002350

Duhen, T., Duhen, R., Montler, R., Moses, J., Moudgil, T., de Miranda, N. F., Goodall, C. P., Blair, T. C., Fox, B. A., McDermott, J. E., et al. (2018). Co-expression of CD39 and CD103 identifies tumor-reactive CD8 T cells in human solid tumors. Nature communications 9, 2724. 10.1038/s41467-018-05072-0

Dusseaux, M., Martin, E., Serriari, N., Peguillet, I., Premel, V., Louis, D., Milder, M., Le Bourhis, L., Soudais, C., Treiner, E., and Lantz, O. (2011). Human MAIT cells are xenobiotic-resistant, tissue-targeted, CD161hi IL-17-secreting T cells. Blood 117, 1250–1259. 10.1182/blood-2010-08-303339

Fornes, O., Castro-Mondragon, J. A., Khan, A., van der Lee, R., Zhang, X., Richmond, P. A., Modi, B. P., Correard, S., Gheorghe, M., Baranasic, D., et al. (2020). JASPAR 2020: update of the openaccess database of transcription factor binding profiles. Nucleic Acids Res 48, D87–D92. 10.1093/nar/gkz1001

Galletti, G., De Simone, G., Mazza, E. M. C., Puccio, S., Mezzanotte, C., Bi, T. M., Davydov, A. N., Metsger, M., Scamardella, E., Alvisi, G., et al. (2020). Two subsets of stem-like CD8(+) memory T cell progenitors with distinct fate commitments in humans. Nat Immunol 21, 1552–1562. 10.1038/s41590-020-0791-5

Getnet, D., Grosso, J. F., Goldberg, M. V., Harris, T. J., Yen, H. R., Bruno, T. C., Durham, N. M., Hipkiss, E. L., Pyle, K. J., Wada, S., et al. (2010). A role for the transcription factor Helios in human CD4(+)CD25(+) regulatory T cells. Mol Immunol 47, 1595–1600. 10.1016/j.molimm.2010.02.001

Goeppert, B., Frauenschuh, L., Zucknick, M., Stenzinger, A., Andrulis, M., Klauschen, F., Joehrens, K., Warth, A., Renner, M., Mehrabi, A., et al. (2013). Prognostic impact of tumour-infiltrating immune cells on biliary tract cancer. Br J Cancer 109, 2665–2674. 10.1038/bjc.2013.610

Goswami, S., Apostolou, I., Zhang, J., Skepner, J., Anandhan, S., Zhang, X., Xiong, L., Trojer, P., Aparicio, A., Subudhi, S. K., et al. (2018). Modulation of EZH2 expression in T cells improves efficacy of anti-CTLA-4 therapy. J Clin Invest 128, 3813–3818. 10.1172/JCI99760

Grant, F. M., Yang, J., Nasrallah, R., Clarke, J., Sadiyah, F., Whiteside, S. K., Imianowski, C. J., Kuo, P., Vardaka, P., Todorov, T., et al. (2020). BACH2 drives quiescence and maintenance of resting Treg cells to promote homeostasis and cancer immunosuppression. J Exp Med 17. 10.1084/jem.20190711

Gros, A., Robbins, P. F., Yao, X., Li, Y. F., Turcotte, S., Tran, E., Wunderlich, J. R., Mixon, A., Farid, S., Dudley, M. E., et al. (2014). PD-1 identifies the patient-specific CD8(+) tumor-reactive repertoire infiltrating human tumors. J Clin Invest 124, 2246–2259. 10.1172/JCI73639

Gubin, M. M., Zhang, X., Schuster, H., Caron, E., Ward, J. P., Noguchi, T., Ivanova, Y., Hundal, J., Arthur, C. D., Krebber, W. J., et al. (2014). Checkpoint blockade cancer immunotherapy targets tumour-specific mutant antigens. Nature 515, 577–581. 10.1038/nature13988

Hafemeister, C., and Satija, R. (2019). Normalization and variance stabilization of single-cell RNA-seq data using regularized negative binomial regression. Genome Biol 20, 296. 10.1186/s13059-019-1874-1

Hill, J. A., Feuerer, M., Tash, K., Haxhinasto, S., Perez, J., Melamed, R., Mathis, D., and Benoist, C. (2007). Foxp3 transcription-factor-dependent and-independent regulation of the regulatory T cell transcriptional signature. Immunity 27, 786–800. 10.1016/j.immuni.2007.09.010

Huber, J. P., and Farrar, J. D. (2011). Regulation of effector and memory T-cell functions by type I interferon. Immunology 132, 466–474. 10.1111/j.1365-2567.2011.03412.x

Huynh-Thu, V. A., Irrthum, A., Wehenkel, L., and Geurts, P. (2010). Inferring regulatory networks from expression data using tree-based methods. PLoS One 5. 10.1371/journal.pone.0012776

Kasper, H. U., Drebber, U., Stippel, D. L., Dienes, H. P., and Gillessen, A. (2009). Liver tumor infiltrating lymphocytes: comparison of hepatocellular and cholangiolar carcinoma. World J Gastroenterol 15, 5053–5057. 10.3748/wjg.15.5053

Kent, W. J., Zweig, A. S., Barber, G., Hinrichs, A. S., and Karolchik, D. (2010). BigWig and BigBed: enabling browsing of large distributed datasets. Bioinformatics 26, 2204–2207. 10.1093/bioinformatics/btq351

Kim, R. D., Chung, V., Alese, O. B., El-Rayes, B. F., Li, D., Al-Toubah, T. E., Schell, M. J., Zhou, J. M., Mahipal, A., Kim, B. H., and Kim, D. W. (2020). A Phase 2 Multi-institutional Study of Nivolumab for Patients With Advanced Refractory Biliary Tract Cancer. JAMA Oncol 6, 888–894. 10.1001/jamaoncol.2020.0930

Kornete, M., Sgouroudis, E., and Piccirillo, C. A. (2012). ICOS-dependent homeostasis and function of Foxp3+ regulatory T cells in islets of nonobese diabetic mice. J Immunol 188, 1064–1074. 10.4049/jimmunol.1101303

Langmead, B., and Salzberg, S. L. (2012). Fast gapped-read alignment with Bowtie 2. Nat Methods 9, 357–359. 10.1038/nmeth.1923

Le, D. T., Uram, J. N., Wang, H., Bartlett, B. R., Kemberling, H., Eyring, A. D., Skora, A. D., Luber, B. S., Azad, N. S., Laheru, D., et al. (2015). PD-1 Blockade in Tumors with Mismatch-Repair Deficiency. N Engl J Med 372, 2509–2520. 10.1056/NEJMoa1500596

Levine, J. H., Simonds, E. F., Bendall, S. C., Davis, K. L., Amir el, A. D., Tadmor, M. D., Litvin, O., Fienberg, H. G., Jager, A., Zunder, E. R., et al. (2015). Data-Driven Phenotypic Dissection of AML Reveals Progenitor-like Cells that Correlate with Prognosis. Cell 162, 184–197. 10.1016/j.cell.2015.05.047

Li, H., Handsaker, B., Wysoker, A., Fennell, T., Ruan, J., Homer, N., Marth, G., Abecasis, G., Durbin, R., and Genome Project Data Processing, S. (2009). The Sequence Alignment/Map format and SAMtools. Bioinformatics 25, 2078–2079. 10.1093/bioinformatics/btp352

Liao, Y., Smyth, G. K., and Shi, W. (2019). The R package Rsubread is easier, faster, cheaper and better for alignment and quantification of RNA sequencing reads. Nucleic Acids Res 47, e47. 10.1093/nar/gkz114

Linderman, G. C., Zhao, J., and Kluger, Y. (2018). Zero-preserving imputation of scRNA-seq data using low-rank approximation. bioRxiv 397588.

Liu, Y., Carlsson, R., Comabella, M., Wang, J., Kosicki, M., Carrion, B., Hasan, M., Wu, X., Montalban, X., Dziegiel, M. H., et al. (2014). FoxA1 directs the lineage and immunosuppressive properties of a novel regulatory T cell population in EAE and MS. Nat Med 20, 272–282. 10.1038/nm.3485

Love, M. I., Huber, W., and Anders, S. (2014). Moderated estimation of fold change and dispersion for RNA-seq data with DESeq2. Genome Biol 15, 550. 10.1186/s13059-014-0550-8

Lugli, E., Galletti, G., Boi, S. K., and Youngblood, B. A. (2020). Stem, Effector, and Hybrid States of Memory CD8(+) T Cells. Trends Immunol 41, 17–28. 10.1016/j.it.2019.11.004

Lugli, E., Zanon, V., Mavilio, D., and Roberto, A. (2017). FACS Analysis of Memory T Lymphocytes. Methods Mol Biol 1514, 31–47. 10.1007/978-1-4939-6548-9_3

Mass, E., Ballesteros, I., Farlik, M., Halbritter, F., Gunther, P., Crozet, L., Jacome-Galarza, C. E., Handler, K., Klughammer, J., Kobayashi, Y., et al. (2016). Specification of tissue-resident macrophages during organogenesis. Science 353. 10.1126/science.aaf4238

Miller, B. C., Sen, D. R., Al Abosy, R., Bi, K., Virkud, Y. V., LaFleur, M. W., Yates, K. B., Lako, A., Felt, K., Naik, G. S., et al. (2019). Subsets of exhausted CD8(+) T cells differentially mediate tumor control and respond to checkpoint blockade. Nat Immunol 20, 326–336. 10.1038/s41590-019-0312-6

Mohamed, J. Y., Faqeih, E., Alsiddiky, A., Alshammari, M. J., Ibrahim, N. A., and Alkuraya, F. S. (2013). Mutations in MEOX1, encoding mesenchyme homeobox 1, cause Klippel-Feil anomaly. Am J Hum Genet 92, 157–161. 10.1016/j.ajhg.2012.11.016

Nakamura, H., Arai, Y., Totoki, Y., Shirota, T., Elzawahry, A., Kato, M., Hama, N., Hosoda, F., Urushidate, T., Ohashi, S., et al. (2015). Genomic spectra of biliary tract cancer. Nat Genet 47, 1003–1010. 10.1038/ng.3375

Okonechnikov, K., Conesa, A., and Garcia-Alcalde, F. (2016). Qualimap 2: advanced multi-sample quality control for high-throughput sequencing data. Bioinformatics 32, 292–294. 10.1093/bioinformatics/btv566

Piha-Paul, S. A., Oh, D. Y., Ueno, M., Malka, D., Chung, H. C., Nagrial, A., Kelley, R. K., Ros, W., Italiano, A., Nakagawa, K., et al. (2020). Efficacy and safety of pembrolizumab for the treatment of advanced biliary cancer: Results from the KEYNOTE-158 and KEYNOTE-028 studies. Int J Cancer 147, 2190–2198. 10.1002/ijc.33013

Plitas, G., Konopacki, C., Wu, K., Bos, P. D., Morrow, M., Putintseva, E. V., Chudakov, D. M., and Rudensky, A. Y. (2016). Regulatory T Cells Exhibit Distinct Features in Human Breast Cancer. Immunity 45, 1122–1134. 10.1016/j.immuni.2016.10.032

Plitas, G., and Rudensky, A. Y. (2020). Regulatory T Cells in Cancer. Annual Review of Cancer Biology 4, 459–477. 10.1146/annurev-cancerbio-030419-033428

Qian, J., Olbrecht, S., Boeckx, B., Vos, H., Laoui, D., Etlioglu, E., Wauters, E., Pomella, V., Verbandt, S., Busschaert, P., et al. (2020). A pan-cancer blueprint of the heterogeneous tumor microenvironment revealed by single-cell profiling. Cell Res 30, 745–762. 10.1038/s41422-020-0355-0

Quinlan, A. R., and Hall, I. M. (2010). BEDTools: a flexible suite of utilities for comparing genomic features. Bioinformatics 26, 841–842. 10.1093/bioinformatics/btq033

Robinson, J. T., Thorvaldsdottir, H., Winckler, W., Guttman, M., Lander, E. S., Getz, G., and Mesirov, J. P. (2011). Integrative genomics viewer. Nat Biotechnol 29, 24–26. 10.1038/nbt.1754

Sade-Feldman, M., Yizhak, K., Bjorgaard, S. L., Ray, J. P., de Boer, C. G., Jenkins, R. W., Lieb, D. J., Chen, J. H., Frederick, D. T., Barzily-Rokni, M., et al. (2018). Defining T cell states associated with response to checkpoint immunotherapy in melanoma. Cell 175, 998–1013 e1020. 10.1016/j.cell.2018.10.038

Saelens, W., Cannoodt, R., Todorov, H., and Saeys, Y. (2019). A comparison of single-cell trajectory inference methods. Nat Biotechnol 37, 547–554. 10.1038/s41587-019-0071-9

Seglen, P. O. (1976). Preparation of isolated rat liver cells. Methods Cell Biol 13, 29–83. 10.1016/s0091-679x(08)61797-5

Sia, D., Hoshida, Y., Villanueva, A., Roayaie, S., Ferrer, J., Tabak, B., Peix, J., Sole, M., Tovar, V., Alsinet, C., et al. (2013). Integrative molecular analysis of intrahepatic cholangiocarcinoma reveals 2 classes that have different outcomes. Gastroenterology 144, 829–840. 10.1053/j.gastro.2013.01.001

Siddiqui, I., Schaeuble, K., Chennupati, V., Fuertes Marraco, S. A., Calderon-Copete, S., Pais Ferreira, D., Carmona, S. J., Scarpellino, L., Gfeller, D., Pradervand, S., et al. (2019). Intratumoral Tcf1(+)PD-1(+)CD8(+) T cells with stem-like properties promote tumor control in response to vaccination and checkpoint blockade immunotherapy. Immunity 50, 195–211 e110. 10.1016/j.immuni.2018.12.021

Simoni, Y., Becht, E., Fehlings, M., Loh, C. Y., Koo, S. L., Teng, K. W. W., Yeong, J. P. S., Nahar, R., Zhang, T., Kared, H., et al. (2018). Bystander CD8(+) T cells are abundant and phenotypically distinct in human tumour infiltrates. Nature 557, 575–579. 10.1038/s41586-018-0130-2

Skuntz, S., Mankoo, B., Nguyen, M. T., Hustert, E., Nakayama, A., Tournier-Lasserve, E., Wright, C. V., Pachnis, V., Bharti, K., and Arnheiter, H. (2009). Lack of the mesodermal homeodomain protein MEOX1 disrupts sclerotome polarity and leads to a remodeling of the cranio-cervical joints of the axial skeleton. Dev Biol 332, 383–395. 10.1016/j.ydbio.2009.06.006

Snyder, A., Makarov, V., Merghoub, T., Yuan, J., Zaretsky, J. M., Desrichard, A., Walsh, L. A., Postow, M. A., Wong, P., Ho, T. S., et al. (2014). Genetic basis for clinical response to CTLA-4 blockade in melanoma. N Engl J Med 371, 2189–2199. 10.1056/NEJMoa1406498

Tariq, N. U., McNamara, M. G., and Valle, J. W. (2019). Biliary tract cancers: current knowledge, clinical candidates and future challenges. Cancer Manag Res 11, 2623–2642. 10.2147/CMAR.S157092

Tirosh, I., Izar, B., Prakadan, S. M., Wadsworth, M. H., 2nd, Treacy, D., Trombetta, J. J., Rotem, A., Rodman, C., Lian, C., Murphy, G., et al. (2016). Dissecting the multicellular ecosystem of metastatic melanoma by single-cell RNA-seq. Science 352, 189–196. 10.1126/science.aad0501

Tumeh, P. C., Harview, C. L., Yearley, J. H., Shintaku, I. P., Taylor, E. J., Robert, L., Chmielowski, B., Spasic, M., Henry, G., Ciobanu, V., et al. (2014). PD-1 blockade induces responses by inhibiting adaptive immune resistance. Nature 515, 568–571. 10.1038/nature13954

Vaine, C. A., and Soberman, R. J. (2014). The CD200-CD200R1 inhibitory signaling pathway: immune regulation and host-pathogen interactions. Adv Immunol 121, 191–211. 10.1016/B978-0-12-800100-4.00005-2

van Dijk, D., Sharma, R., Nainys, J., Yim, K., Kathail, P., Carr, A. J., Burdziak, C., Moon, K. R., Chaffer, C. L., Pattabiraman, D., et al. (2018). Recovering Gene Interactions from Single-Cell Data Using Data Diffusion. Cell 174, 716–729 e727. 10.1016/j.cell.2018.05.061

Vasanthakumar, A., Liao, Y., Teh, P., Pascutti, M. F., Oja, A. E., Garnham, A. L., Gloury, R., Tempany, J. C., Sidwell, T., Cuadrado, E., et al. (2017). The TNF Receptor Superfamily-NF-kappaB Axis Is Critical to Maintain Effector Regulatory T Cells in Lymphoid and Non-lymphoid Tissues. Cell reports 20, 2906–2920. 10.1016/j.celrep.2017.08.068

Vento-Tormo, R., Efremova, M., Botting, R. A., Turco, M. Y., Vento-Tormo, M., Meyer, K. B., Park, J. E., Stephenson, E., Polanski, K., Goncalves, A., et al. (2018). Single-cell reconstruction of the early maternal-fetal interface in humans. Nature 563, 347–353. 10.1038/s41586-018-0698-6

Wang, D., Quiros, J., Mahuron, K., Pai, C. C., Ranzani, V., Young, A., Silveria, S., Harwin, T., Abnousian, A., Pagani, M., et al. (2018). Targeting EZH2 Reprograms Intratumoral Regulatory T Cells to Enhance Cancer Immunity. Cell reports 23, 3262–3274. 10.1016/j.celrep.2018.05.050

Weiskopf, K. (2017). Cancer immunotherapy targeting the CD47/SIRPalpha axis. Eur J Cancer 76, 100–109. 10.1016/j.ejca.2017.02.013

Wolf, F. A., Angerer, P., and Theis, F. J. (2018). SCANPY: large-scale single-cell gene expression data analysis. Genome Biol 19, 15. 10.1186/s13059-017-1382-0

Zappia, L., and Oshlack, A. (2018). Clustering trees: a visualization for evaluating clusterings at multiple resolutions. Gigascience 7. 10.1093/gigascience/giy083

Zhang, L., Li, Z., Skrzypczynska, K. M., Fang, Q., Zhang, W., O’Brien, S. A., He, Y., Wang, L., Zhang, Q., Kim, A., et al. (2020a). Single-Cell Analyses Inform Mechanisms of Myeloid-Targeted Therapies in Colon Cancer. Cell 181, 442–459 e429. 10.1016/j.cell.2020.03.048

Zhang, M., Yang, H., Wan, L., Wang, Z., Wang, H., Ge, C., Liu, Y., Hao, Y., Zhang, D., Shi, G., et al. (2020b). Single-cell transcriptomic architecture and intercellular crosstalk of human intrahepatic cholangiocarcinoma. J Hepatol 73, 1118–1130. 10.1016/j.jhep.2020.05.039

Zhang, Y., Liu, T., Meyer, C. A., Eeckhoute, J., Johnson, D. S., Bernstein, B. E., Nusbaum, C., Myers, R. M., Brown, M., Li, W., and Liu, X. S. (2008). Model-based analysis of ChIP-Seq (MACS). Genome Biol 9, R137. 10.1186/gb-2008-9-9-r137

Zheng, G. X., Terry, J. M., Belgrader, P., Ryvkin, P., Bent, Z. W., Wilson, R., Ziraldo, S. B., Wheeler, T. D., McDermott, G. P., Zhu, J., et al. (2017). Massively parallel digital transcriptional profiling of single cells. Nature communications 8, 14049. 10.1038/ncomms14049

