## Supplementary material for "Multimodal single-cell profiling of intrahepatic cholangiocarcinoma defines hyperactivated Tregs as a potential therapeutic target"

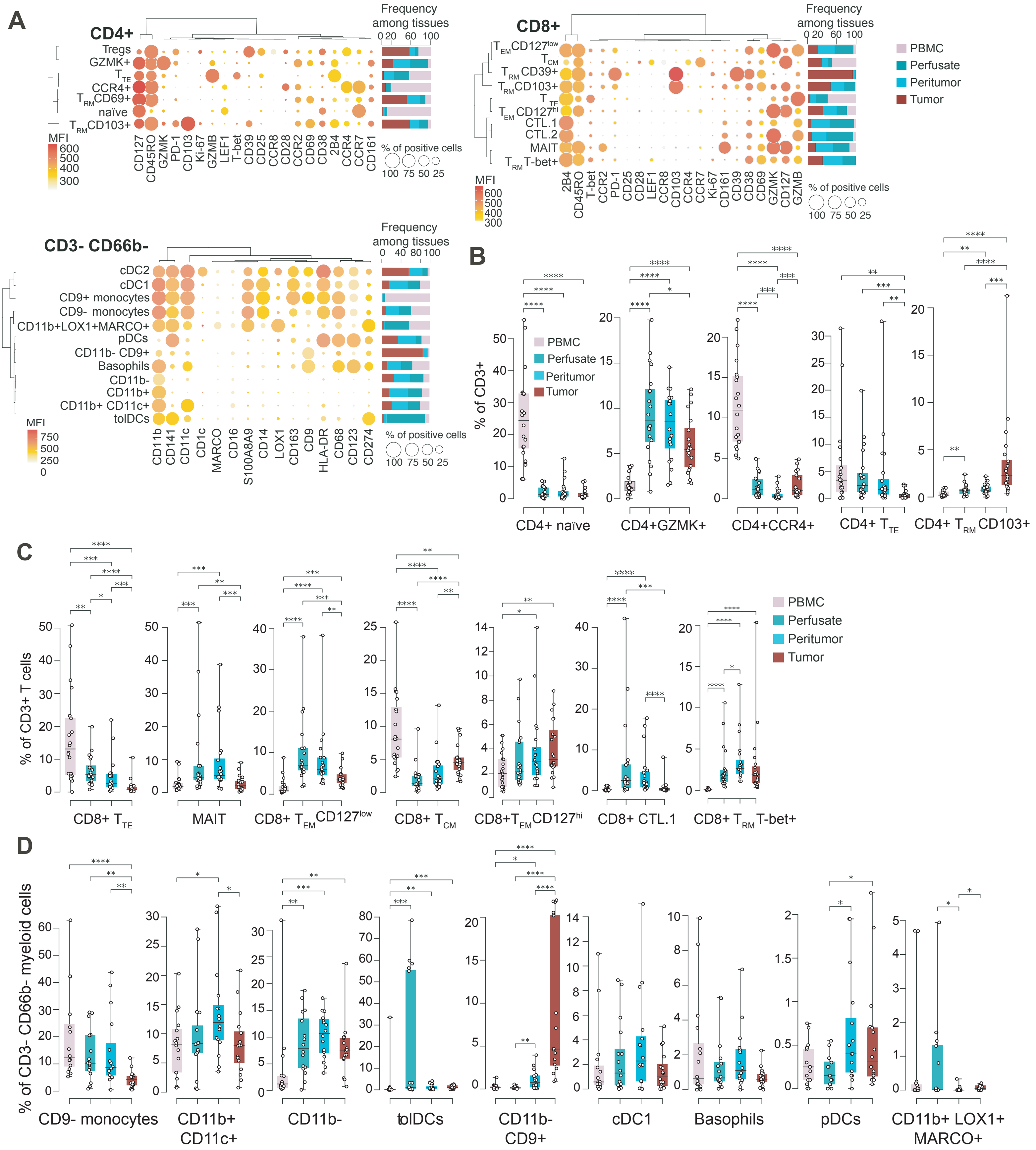


**Figure S1. T cell and myeloid cell landscapes of intrahepatic cholangiocarcinoma. A.** Balloon plot maps showing the percent expression and the mean fluorescent intensity (MFI) of specific markers (columns) in discrete PhenoGraph clusters (rows) of CD4+ and CD8+ T cells, and of CD3- CD66b- myeloid cells. Hierarchical meta-clustering grouped markers and clusters with similar immunophenotypes. Bar plots show the percent frequency of each cluster among different tissues. **B-D.** Box plots showing the median and the IQR of PhenoGraph cluster frequency at different tissue sites. Bars indicate the SD. Dots depict values of single patients. *=P<0.05, **=P<0.01, ***=P<0.001; ****=P<0.0001, two-sided Mann-Whitney test.

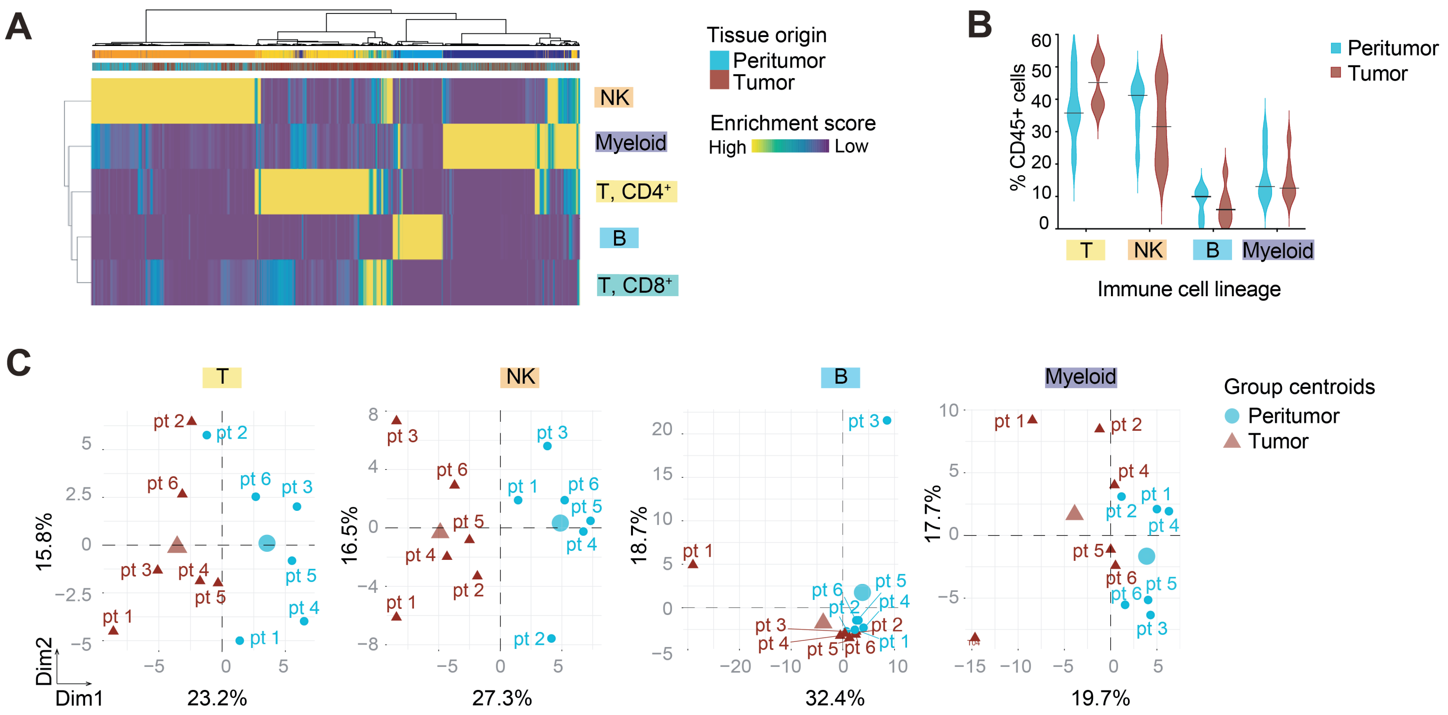


**Figure S2. The landscape of immune infiltrates in human iCCA revealed by scRNA-seq. A.** Heatmap of SingleR enrichment scores of single CD45+ cells (column) to each "Main Immune Cell Expression Data'" reference signature (row) after Adaptively-threshold Low-Rank Approximation (ALRA) imputation. Cells were further meta-clustered based on SingleR enrichment scores. **B.** Violin plots showing the relative frequency of immune cell types identified in A. Lines represent median frequencies (n=6 patients). **C.** PCA plots showing the distribution of peritumoral (light blue circles) and tumoral (dark red triangles) samples from 6 patients (pt) according the gene expression profile of each sample. Symbols of bigger size indicate the centroid of the distribution.

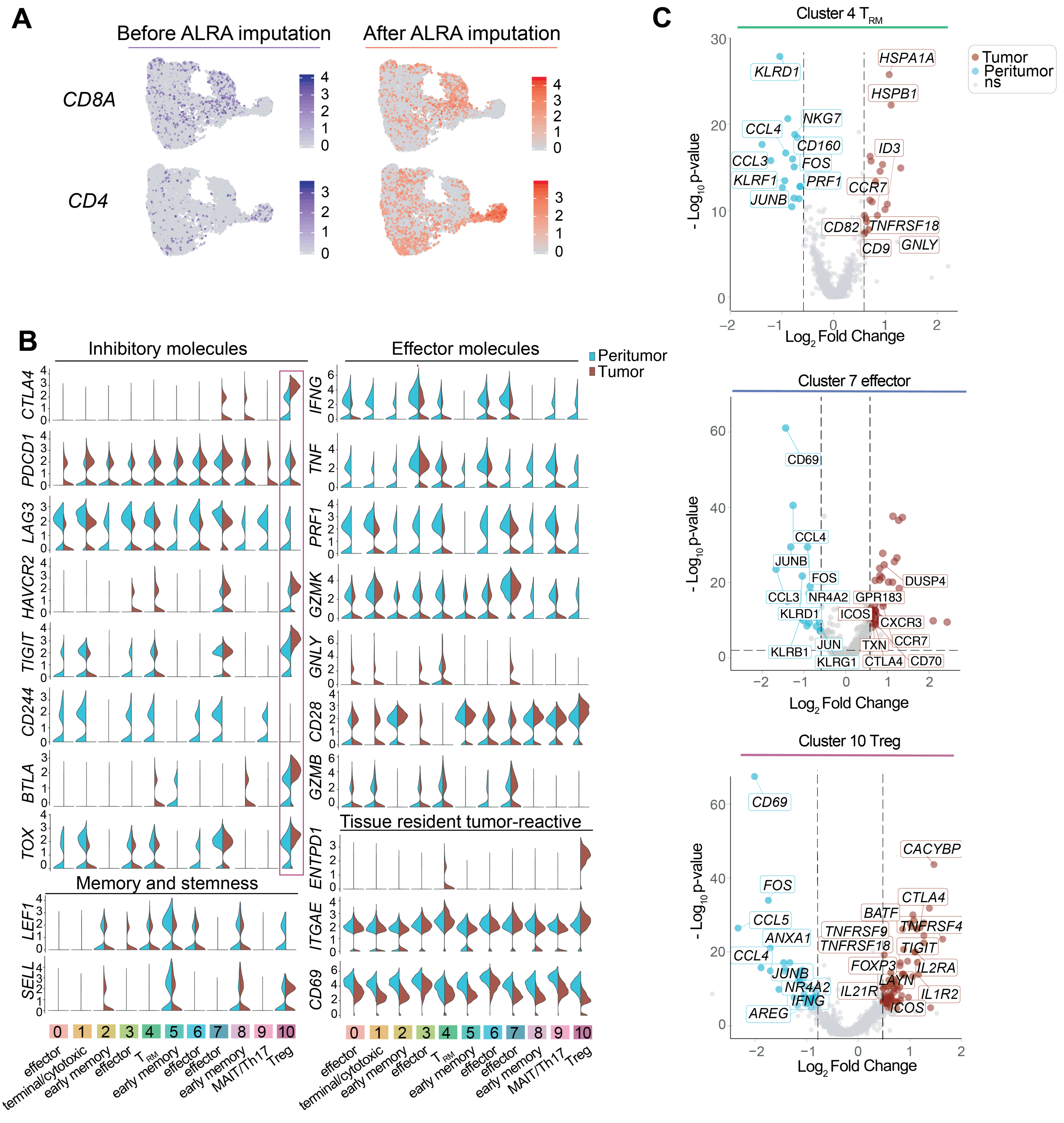


**Figure S3. The identity of iCCA-infiltrating cells as revealed by scRNA-seq. A.** UMAP plots showing the expression of *CD4* and *CD8A* genes before and after Adaptively-threshold Low-Rank Approximation (ALRA) imputation. **B.** Expression of functionally relevant genes in T cell clusters. Width of the violin plot denotes average expression levels. **C.** Volcano plot showing differentially expressed genes between tumoral (dark red) and peritumoral (light blue) cells from scRNA-seq cluster 4 ($T_{RM}$), cluster 7 (effector) and cluster 10 (Tregs); (q-value<0.01; |FC|>1.5)

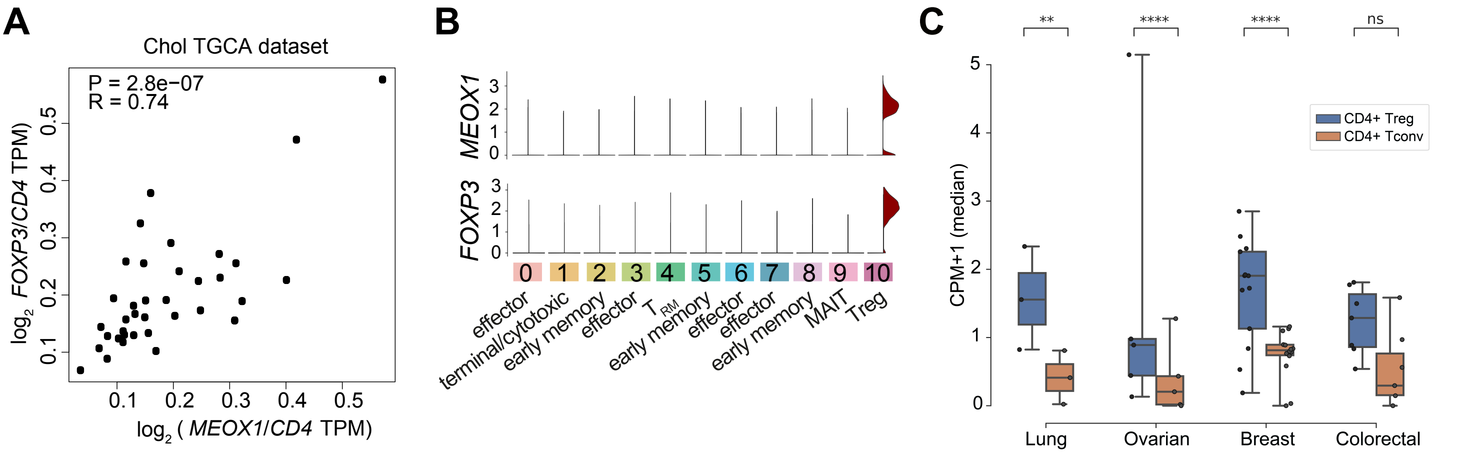


**Figure S4. The gene expression profile of Tregs infiltrating human cholangiocarcinoma. A.** Pair-wise correlation between *MEOX1* and *FOXP3* expression in the The Cancer Genoma Atlas (TGCA) cholangiocarcinoma (Chol) dataset (n=36), as from Gene Expression Profiling Interactive Analysis (GEPIA2). *MEOX1* and *FOXP3* expression have been normalized by *CD4* expression. P: p-value; R: Pearson correlation coefficient. **B.** Imputed expression levels of *MEOX1* and *FOXP3* among single-cell RNA-seq T cell clusters identified in Fig. 3. Violin plot's colours denote tissue of origin: tumoral (dark red) and peritumoral (light blue); widths denote average expression levels. **C.** Heatmap depicting *MEOX1* expression levels in Treg and CD4+ conventional T cells (Tconv) infiltrating lung (n=3), ovarian (n=5), breast (n=14) and colorectal (n=7) tumors, as from Qian et al, Cell Res, 2020 (PMID: 32561858). **=P<0.01, ****=P<0.0001, Wilcoxon test. ns: not significant.
